# Virus-induced paraspeckle-like condensates are essential hubs for gene expression and their formation drives genomic instability

**DOI:** 10.1101/2024.07.23.604779

**Authors:** Katherine L. Harper, Elena Harrington, Connor Hayward, Wiyada Wongwiwat, Robert E White, Adrian Whitehouse

**Author notes:** Correspondence: Adrian Whitehouse. Authors contributed equally (Katherine L. Harper, Elena Harrington).

## Abstract

The nucleus is a highly structured environment containing multiple membrane-less bodies formed through liquid-liquid phase separation. These provide spatial separation and concentration of specific biomolecules enabling efficient and discrete processes to occur which regulate gene expression. One such nuclear body, paraspeckles, are comprised of multiple paraspeckle proteins (PSPs) built around the architectural RNA, *NEAT1_2*. Paraspeckle function is yet to be fully elucidated but has been implicated in a variety of developmental and disease scenarios. We demonstrate that Kaposi’s sarcoma-associated herpesvirus (KSHV) drives formation of structurally distinct paraspeckles with a dramatically increased size and altered protein composition that are essential for productive lytic replication. We highlight these virus-induced paraspeckle-like structures form adjacent to virus replication centres, functioning as RNA processing hubs for both viral and cellular transcripts during infection. Notably, we reveal that PSP sequestration into virus-induced paraspeckle-like structures results in increased genome instability during both KSHV and Epstein Barr virus (EBV) infection, implicating their formation in virus-mediated tumorigenesis.

## Introduction

The nucleus is an extremely crowded but highly organised environment, containing several distinct nuclear bodies which each fulfil specific roles and functions. These membrane-less organelles are formed through liquid-liquid phase separation (LLPS), allowing spatial separation and concentration of specific components enabling efficient discrete processes to regulate gene expression (1). Paraspeckles, discovered relatively recently in 2002, are one such nuclear body, located in the interchromatin space (2). Canonical paraspeckles form through core paraspeckle protein (PSP) interactions with the architectural lncRNA *NEAT1_2,* producing a unique structure comprising of an outer shell and inner core (3). Biogenesis occurs co-transcriptionally, with rapid binding of the core PSPs SFPQ and NONO aiding in *NEAT1_2* stability. Oligomerisation of these PSPs along the ∼23kb transcript leads to formation of a stable ribonucleoprotein particle (RNP) (4, 5). LLPS occurs as multiple RNPs are then linked together through FUS, leading to the formation of mature paraspeckles (6). There are estimated to be around 100 proteins that dynamically associate with paraspeckles, categorised as either essential, important or dispensable for paraspeckle formation (5, 7). The majority of mammalian tissues and cells contain paraspeckles, however depending on cell type paraspeckles can vary in number, ranging between 5 and 20 foci per nucleus (8). Paraspeckle function has yet to be fully determined but they have been implicated in a range of cellular processes, such as regulation of gene expression by sequestration of RNA and proteins which, when dysregulated, lead to various disease scenarios. This role complements their dynamic structure, with their formation, function and dissolution orchestrating a fine-tuned cellular response to specific stimuli (9–12).

Kaposi’s sarcoma-associated herpesvirus (KSHV) is a large dsDNA gammaherpesvirus associated with the development of Kaposi’s sarcoma (KS), a highly vascular tumour of endothelial lymphatic origin, and several other AIDs-associated malignancies including primary effusion lymphoma (PEL) and some forms of multicentric Castleman’s disease (13–17). KSHV exhibits a biphasic life cycle consisting of latent persistence and lytic replication. Latency is established in B cells and in the tumour setting, where viral gene expression is limited to the latency-associated nuclear antigen (LANA), viral FLICE inhibitory protein, viral cyclin, kaposins and several virally-encoded miRNAs (18–20). Upon reactivation, KSHV enters the lytic replication phase, leading to the highly orchestrated expression of more than 80 viral proteins that are sufficient to produce infectious virions (21, 22). In KS lesions, most infected cells harbour the virus in a latent state. However, a small proportion of cells undergo lytic and abortive lytic replication that leads to the secretion of angiogenic, inflammatory and proliferative factors that act in a paracrine manner on latently-infected cells to enhance tumorigenesis (23). Lytic replication also leads to genomic instability (24) and sustains KSHV episomes in latently-infected cells that would otherwise be lost during cell division (25, 26). Therefore, both the latent and lytic replication phases are essential for KSHV-mediated tumourigenicity, and the ability to inhibit the lytic replication phase represents a therapeutic intervention strategy for the treatment of KSHV-associated diseases (27, 28).

During the early stages of herpesvirus lytic replication, the nuclear architecture of the host cell undergoes a striking re-organization to facilitate viral replication. This is driven by the formation of virus-induced nuclear structures, termed virus replication and transcription compartments (vRTCs), which support viral transcription, DNA replication and packaging, and capsid assembly (29, 30). As lytic replication proceeds, small vRTCs coalesce into a singular large globular structure that ultimately fills most of the nuclear space compressing and marginalising the cellular chromatin to the nuclear periphery. How this drastic virus-induced remodelling of the nuclear architecture affects the localisation and function of nuclear bodies to regulate gene expression is yet to be fully elucidated. In particular, how paraspeckles are impacted is of prominent interest as their dynamic formation is emerging as a global sensor of cellular stress. Notably, multiple stressors including mitochondrial stress, hypoxia and heat shock can induce changes in paraspeckle abundance, which is thought to have a cytoprotective role in response to these stressors (31). This study demonstrates that KSHV drives formation of novel, non-canonical paraspeckles during lytic replication. Specifically, we show for the first time that paraspeckles are altered in their localisation, aligning adjacent to vRTCs, and are approximately 10 times larger than the size of standard paraspeckles. Further evidence supports that these structures are not true canonical paraspeckles, confirmed by their unique protein composition including the association of the KSHV-encoded ORF11 protein which appears to drive puncta biogenesis during infection. Notably, these KSHV-induced paraspeckle-like condensates are essential for virus replication and infectious virion production, acting as hubs for the processing of both cellular and viral RNAs required for efficient KSHV lytic replication. Interestingly, the formation of these virus-induced paraspeckles are herpesvirus subfamily specific, as they are also observed during the lytic replication cycle of the human gamma herpesvirus Epstein Barr virus (EBV), but fail to form during human alpha or beta herpesvirus lytic replication. Importantly, aligned with the unique gamma herpesvirus specificity, a correlation is observed between the formation of paraspeckle-like condensates and increased genomic instability which may indirectly play a role in gamma herpesvirus-mediated oncogenesis (27). Taken together, this study has identified novel virus-induced paraspeckle-like condensates forming during KSHV infection which enhance lytic replication and are implicated in virus-mediated genomic instability.

## Results

### KSHV induces the formation of novel puncta during lytic replication

The host cell nuclear architecture undergoes dramatic remodelling to facilitate KSHV lytic replication, due to the formation of vRTCs which support viral transcription, DNA synthesis and capsid assembly. We therefore aimed to investigate how this reorganisation affects nuclear bodies, with particular focus on the impact of virus-induced remodelling of paraspeckles, a nuclear body whose formation is emerging as a global sensor of cellular stress (9, 31). Interestingly, other cell stress sensors such as cytoplasmic stress granules are disrupted during KSHV lytic replication (32). Therefore, to examine whether KSHV-mediated remodelling of the nucleus induces changes in paraspeckles, TREx-BCBL1-RTA cells, a KSHV-latently infected B-lymphocyte cell line containing a Myc-tagged version of the viral RTA under the control of a doxycycline-inducible promoter were utilised, allowing efficient induction of the lytic cascade with addition of doxycycline. Cells remained latent or reactivated over a time course prior to immunostaining with antibodies against SFPQ and RNA pol II, markers of paraspeckles and vRTCs, respectively. vRTC development was observed from 8 hours post lytic reactivation and reached full maturity by 24 hours, indicated by distinct restructuring of RNA pol II into an elliptical configuration (**Fig. 1Ai-ii**). Surprisingly, a striking change was observed upon staining of the paraspeckle marker, SFPQ. Notably, in KSHV-latently infected TREx-BCBL1-RTA cells, no paraspeckles were observed and SFPQ remained diffuse throughout the nucleus. Whether paraspeckles exist in non-KSHV infected B cells is currently unknown, however paraspeckle formation in response to stress has been reported in many but not all cultured cells, with large amounts of cell type variation (33, 34). In contrast, during early stages of KSHV lytic replication SFPQ coalesced into distinct large puncta. Whilst these puncta were spatially distinct to nuclear speckles and vRTCs, these SFPQ puncta localised adjacent and around vRTCs, suggesting they may have a complementary function (**Fig. 1Ai-ii, Fig.S1A**). To investigate whether these puncta were specifically induced by KSHV lytic replication, latently-infected TREx-BCBL1-RTA cells were exposed to a number of stress treatments such as serum starving and osmotic stress, with several of these stress treatments known to induce canonical paraspeckle formation (9, 35). Results confirmed SFPQ puncta formation was a virus-specific response, with no puncta observed in latently-infected stress-treated cells (**Fig.S1Bi-ii**). This was further corroborated by formation of SFPQ puncta during lytic replication in an additional KSHV-infected cell line, HEK-293T-rKSHV.219s (**Fig.S1C**), with no puncta observed in latent cells or uninfected HEK-293Ts (**Fig.S1D)**. Together these results suggest that KSHV induces the formation of SFPQ containing puncta during the early stages of its lytic replication cycle.

**Figure 1:**
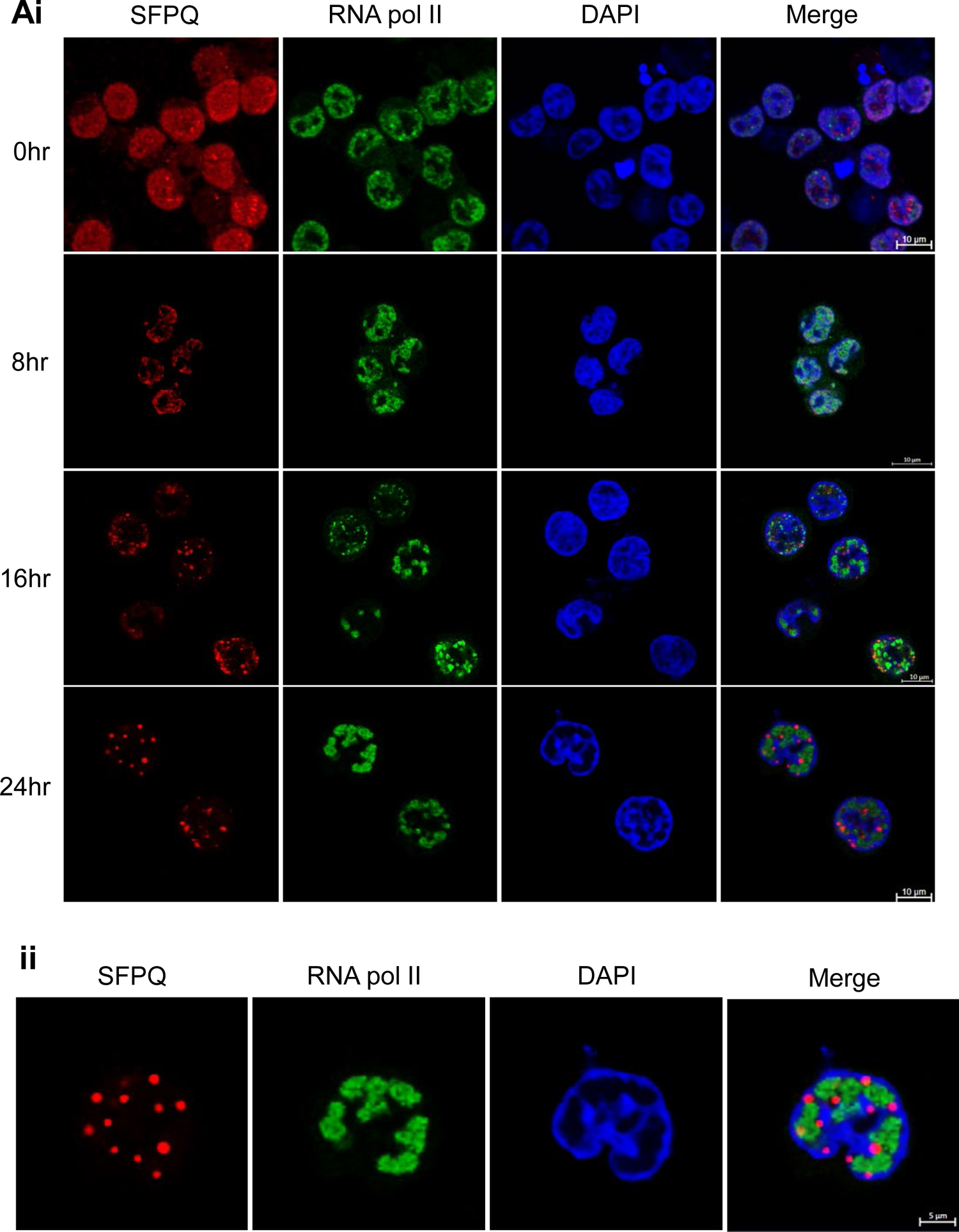
SFPQ forms puncta during gammaherpesvirus lytic replication. Ai: IF of TREx cells at 0, 8, 16 and 24 hours post-reactivation stained against SFPQ (red) and RNA pol II (green). Aii: Greater magnification view of 24hr TREx with antibodies against RNA pol II (green) and SFPQ (red). Scale bars represent 10 µm in Ai and 5 µm in Aii

### KSHV-induced SFPQ containing puncta are paraspeckle-like structures

SFPQ is a multi-functional RBP with numerous roles in cell regulation, including transcriptional repression, alternative splicing and DNA damage. Critically, SFPQ is a core paraspeckle specific protein (PSP) (36). Therefore, to determine whether these novel KSHV-induced puncta only contained aggregated SFPQ protein or were modified paraspeckles, RNA-FISH and immunofluorescence studies were performed to determine whether other paraspeckle components were also contained in these novel puncta. Canonical paraspeckles have a sub-structure comprising core and shell zones, containing different PSPs built around the architectural RNA, *NEAT1_2* (later referred to as just *NEAT1*). RNA FISH confirmed the presence and co-localisation of *NEAT1* in the virus-induced puncta (**Fig. 2A**). Immunostaining with antibodies against additional characterised PSPs also confirmed PSPC1 and NONO co-localise to the puncta (**Fig.2B, Fig.S2A**). This was further supported by co-immunoprecipitation studies showing SFPQ interacts with NONO and PSPC1 during both latency and lytic replication, due to the natural affinity of SFPQ to heterodimerise with these proteins (**Fig.S2B**) (37). Surprisingly however, certain core paraspeckle proteins were missing from the virus-induced paraspeckles. FUS, a core PSP, essential for mature paraspeckle formation is absent from the virus-induced paraspeckles, instead localising to vRTCs. This relocalisation was confirmed by co-immunoprecipitation assays which show a decrease in the association of SFPQ and FUS during KSHV lytic replication compared to latent cells (**Fig.S2C-D**). Furthermore, to examine whether FUS was required for formation of these novel puncta, stable TREx-BCBL1-RTA cells were produced depleting FUS with targeted shRNAs (**Fig.S2E-F**). Results clearly show that paraspeckle-like condensates were still capable of forming in the absence of FUS (**Fig.2C**).

**Figure 2:**
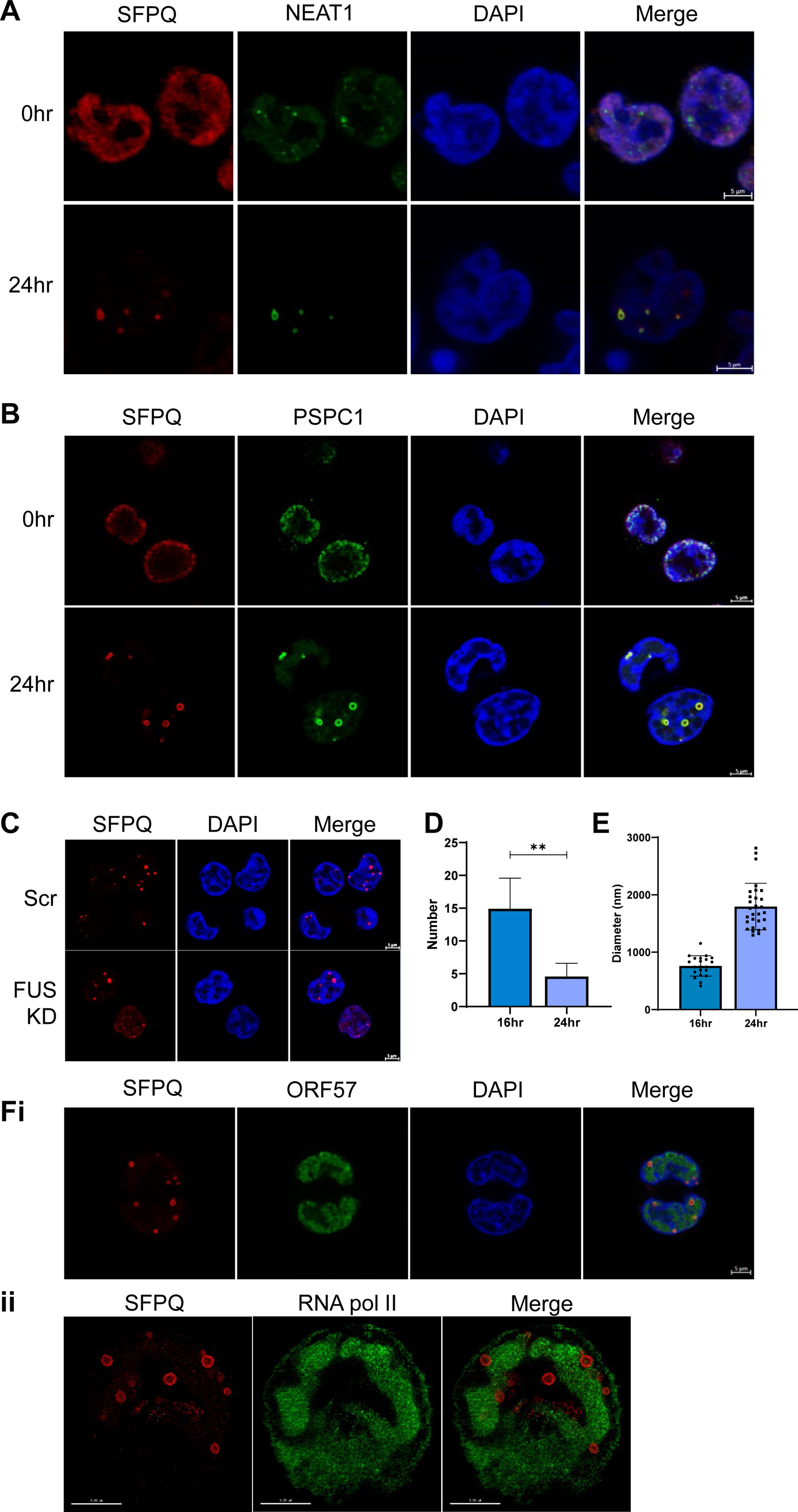
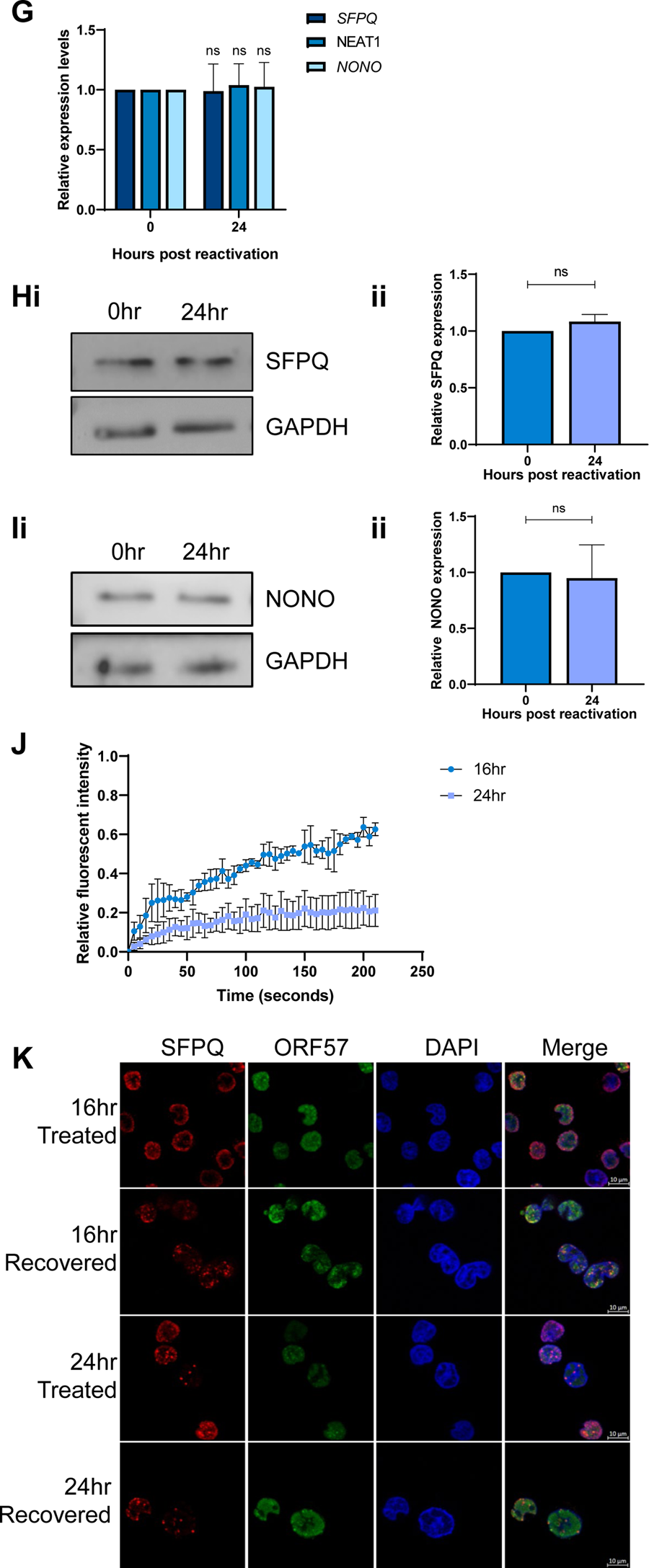
Puncta are characterised as virus-induced paraspeckle-like condensates. IF of TREx-BCBL1-RTA cells at 0 and 24 hours stained for SFPQ (red) and NEAT1 (green) (A) or SFPQ (red) and PSPC1 (green) (B). (C). IF of scr and FUS KD TREx at 24 hours with staining for SFPQ (red). (D). Average number of puncta per z-frame puncta in TREx-BCBL1-RTA. 20 individual cells were analysed via Zen Blue and FIJI software for each time point. (E). Average diameter of puncta in TREx-BCBL1-RTA cells with 16hr or 24 hours post lytic KSHV induction as measured through Zen blue software. (F). IF of SFPQ (red) and ORF57 (green) in 24 hours reactivated TREx-BCBL1-RTA cells. ii STED of SFPQ (red) and RNA pol II (green) at 24 hours in TREx-BCBL1-RTA cells. (G). qPCR analysis of NEAT1, SFPQ and NONO levels in TREx-BCBL1-RTA cells with latent KSHV or 24 hours post lytic KSHV induction, with GAPDH used as a housekeeper (n=3). (H-I) Western blot and densitometry analysis of SFPQ and NONO levels in TREx-BCBL1-RTA cells with latent KSHV or 24 hours post lytic KSHV induction. GAPDH was used as a loading control (n=3). (J). FRAP analysis of puncta at 16 and 24 hours post KSHV lytic induction, measuring fluorescent recovery in TREx-BCBL1-RTA SFPQ-GFP OE cells (n=3). (K). IF of TREx-BCBL1-RTA cells at 16 and 24 hours post lytic induction, stained for SFPQ (red) and ORF57 (green). Cells were treated with 1,6HD and either fixed immediately or allowed to recover. All scale bars are 5 µm except for 10 µm in K. In D-E, G-J data are presented as mean ± SD. Unpaired student T-test was used. All repeats are biological. *P < 0.05, **P < 0.01 and ***P < 0.001.

This analysis suggests there are distinct differences in the formation and structure of the virus-induced paraspeckles compared to canonical paraspeckles. Therefore, to further investigate the formation of KSHV-induced paraspeckle-like structures, TREx-BCBL1-RTA cells were visualised over a lytic replication time course post-reactivation. Paraspeckle-like puncta formation is observed from as early as 8 hours post-reactivation, with the majority forming between 16 and 24 hours (**Fig.S2G**). During this crucial period, as puncta reduced in number, from an average of 15 to 5 per central Z frame slice (**Fig.2D**) they concurrently grew in size, significantly increasing from an average of ∼700 nm to ∼1800 nm, with the largest up to 3000 nm in diameter, which is 10x larger than canonical paraspeckles (**Fig.2E**). This was further supported by high-resolution Airy-Scan microscopy (**Fig.2Fi**), which showed that SFPQ formed a distinct ring structure. Whilst SFPQ predominantly localises to the edge of the ring structure, STED microscopy confirmed it is also distributed throughout the puncta. Furthermore, the higher resolution microscopy suggests a more complex and heterogeneous order on the edge of the rings, as identified by the uneven SFPQ distribution on the periphery **(Fig.2Fii)** thus eluding to a higher structure within the condensates.

Paraspeckle biogenesis occurs co-transcriptionally with core paraspeckle proteins binding to and stabilising *NEAT1*, as such *NEAT1* levels directly correlate with paraspeckle size [7]. Due to the dramatically increased size of the KSHV-induced paraspeckle-like puncta, it was hypothesised that *NEAT1* levels would increase during lytic replication. However, surprisingly, qPCR analysis comparing *NEAT1* levels in latent and reactivated TREx-BCBL1-RTA cells, showed no increase in *NEAT1* expression (**Fig.2G**). Furthermore, qPCR and immunoblot analysis of SFPQ and NONO also showed no significant increase in mRNA (**Fig.2G**) or protein levels during lytic replication (**Fig.2H-I**). Together these data suggest the puncta are non-canonical paraspeckles, therefore we now refer to them as novel virus-induced paraspeckle-like structures.

### KSHV-induced paraspeckles-like structures are dynamic condensates

The dynamic nature of KSHV-induced paraspeckle-like puncta formation suggested they may be formed through liquid-liquid phase separation, a property typically associated with condensates. As such, fluorescent recovery after photobleaching (FRAP) was performed using a TREx-BCBL1-RTA cell line over-expressing GFP-SFPQ. Puncta formed at both 16 and 24 hours post-lytic replication were exposed to photobleaching to probe their potential dynamicity. Significant fluorescent recovery was observed for puncta formed 16 hours post reactivation, whereas puncta formed after 24 hours of lytic replication showed little recovery (**Fig.2J, Fig.S2H**). This is suggestive of a shift in the material state of the paraspeckle-like puncta from liquid-like to gel-like, implicating condensate maturation. Further studies utilised the aliphatic alcohol 1,6-hexanediol, which is capable of dissolving liquid-like states, in contrast, gel-like states are more resistant (38). Similar to the FRAP analysis, puncta formed 16 hours post KSHV lytic reactivation were susceptible to dissolving, whereas puncta formed later at 24 hours post induction were resistant to 1,6-hexanediol treatment. Finally, allowing a 30 minute recovery period post 1,6-hexanediol treatment led to the reforming of puncta at 16 hours post lytic induction, confirming their dynamic liquid-like structure during the early stages of KSHV lytic replication (**Fig.2K**). Conversely, the recovery period failed to increase the number of puncta observed after 24 hours post reactivation, further suggesting these puncta are fully mature and have entered a gel-like state at later stages of replication. Taken together, these results highlight that KSHV-induced paraspeckle-like structures show distinct properties associated with condensates.

### KSHV ORF11 protein drives formation of virus-induced paraspeckle-like condensates

Previous results highlight distinct differences in the structure and composition of KSHV-induced paraspeckles compared to canonical paraspeckles. To analyse these differences in more detail, the proteome composition of virus-induced paraspeckles was compared to canonical paraspeckles and the presence of any KSHV-encoded proteins in these virus-induced structures was interrogated. As previously highlighted, immunofluorescence studies show SFPQ is completely redistributed from a diffuse nuclear staining into KSHV-induced paraspeckle-like condensates at 24 hours post reactivation (**Fig.1A**), therefore affinity pulldowns were performed using anti-SFPQ or control anti-IgG antibodies in latent or 24 hour reactivated KSHV-infected cells, coupled with TMT-labelled quantitative mass spectrometry. Negative control (anti-IgG pulldown) protein abundance was subtracted from total abundance for each protein followed by a fold change comparison between KSHV latent and lytic samples. As expected, canonical paraspeckle proteins were observed in the SFPQ interaction profile during lytic replication (**Fig.S3Ai-iii**). However, interestingly, a selection of DEAD/DEAH box helicases, hnRNPs and the viral ORF11 protein were also identified as interactors with SFPQ exclusively during lytic replication (**Fig.3A**). These are all novel SFPQ interactors and results suggest these proteins are recruited into the virus-induced paraspeckle-like structures and may play a role in condensate function during lytic replication.

**Figure 3:**
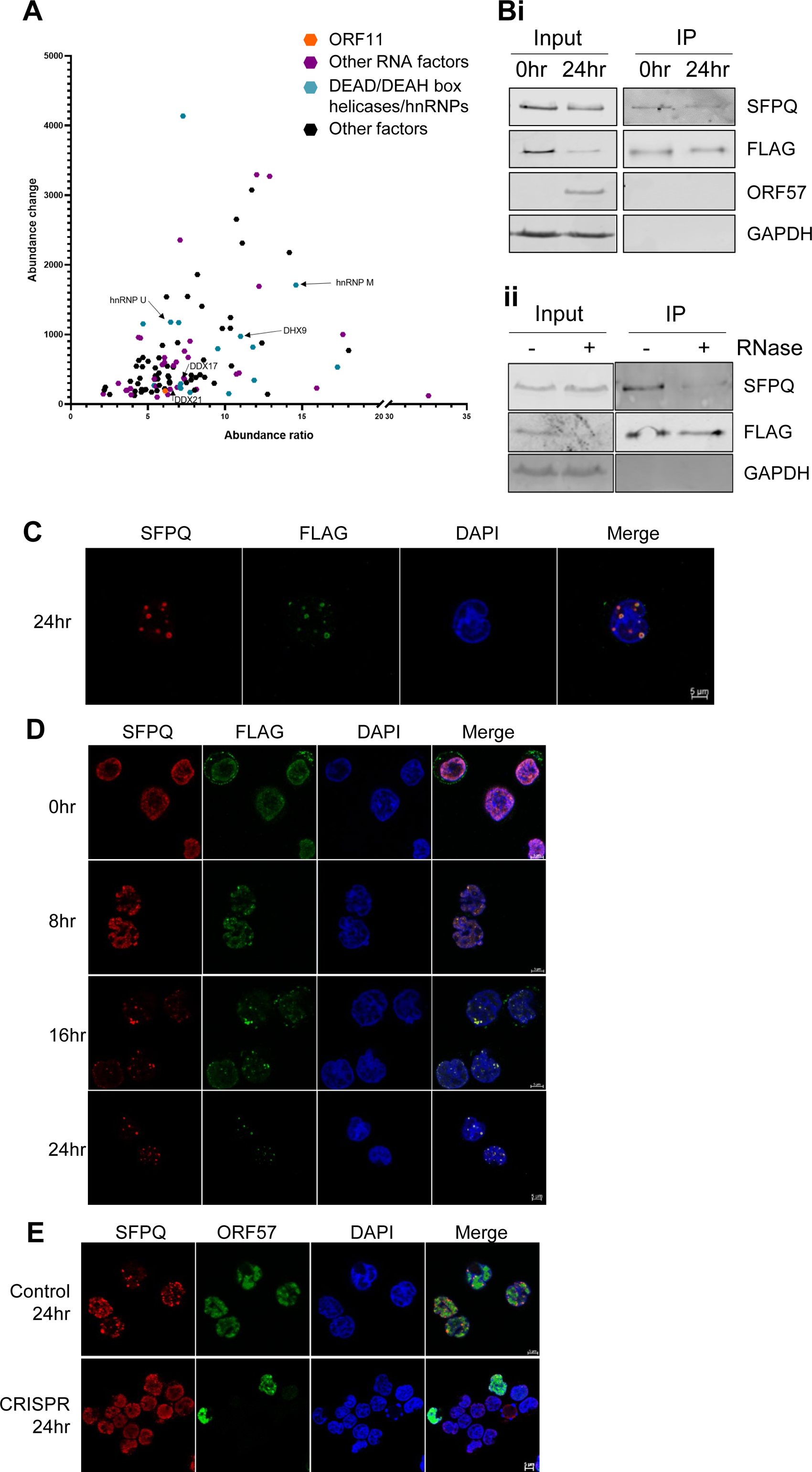
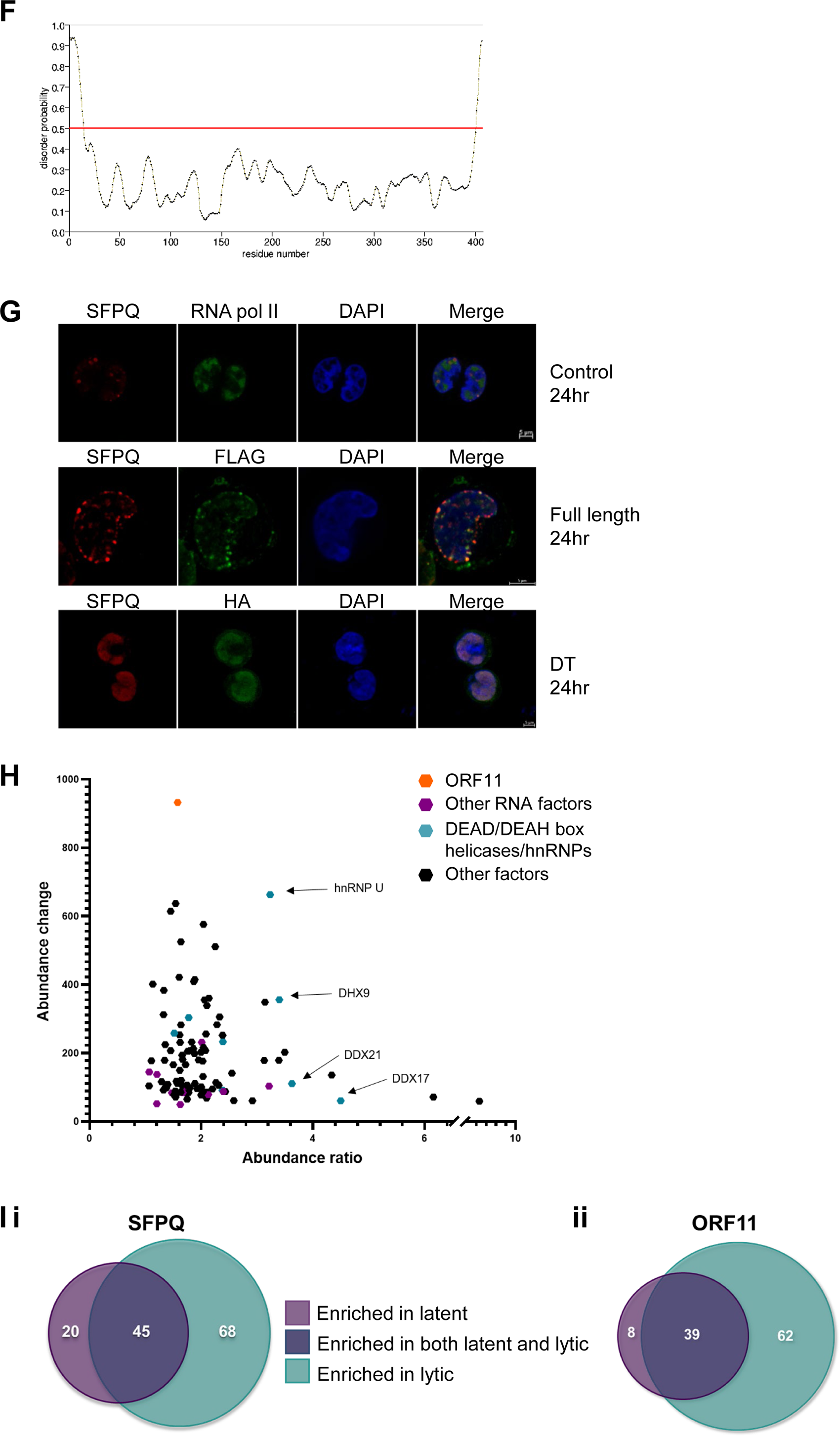
ORF11 is the viral driver in condensate formation. (A). Graph of proteins over a 10% abundance ratio and 4% abundance change in the SFPQ TMT-MS at 24 hours (n=2). ORF11 is highlighted in orange, proteins involved in RNA processing are highlighted purple and DEAD/DEAH box helicases and hnRNP proteins are blue. Proteins confirmed via IF as co-localising are marked with arrows. (B). (i) Western blot analysis of FLAG Co-IPs in TREx FLAG-ORF11 O/E cells harbouring KSHV during latent or 24 hour post lytic replication and probed with antibodies against FLAG (ORF11), SFPQ, ORF57 and GAPDH (n=3). (ii). Western blot analysis of FLAG Co-IPs in TREx FLAG-ORF11 O/E cells at 24 hours +/-RNase. Probed with antibodies against FLAG (ORF11), SFPQ and GAPDH. (C). IF for SFPQ (red) and FLAG (ORF11) (green) after 24 hours of lytic KSHV replication in TREx FLAG-ORF11 O/E cells. (D). IF of FLAG(ORF11) (green) and SFPQ (red) at 0, 8, 16 and 24 hours in TRExFLAG-ORF11 O/E cells harbouring latent KSHV or 8, 16, and 24 hours post lytic induction. (E). IF staining for ORF57(green) and SFPQ (red) in Control and ORF11 CRISPR TREx cells during latent KSHV infection or after 24 hours lytic replication. (F). Protein disorder prediction for ORF11 using PrDOS software. (G). IF in control TREx cells, and rescues of either full length FLAG-ORF11 or double truncated (DT) ORF11-HA in ORF11 CRISPR cells. Staining for SFPQ (red), and either RNA pol II (green), FLAG (green) or HA (green). (H). Graph of proteins over a 10% abundance ratio and 4% abundance enriched in the FLAG-ORF11 TMT-MS at 24 hours (n=2). ORF11 is highlighted in orange, proteins involved in RNA processing are highlighted purple and DEAD/DEAH box helicases and hnRNP proteins are blue. Proteins confirmed via IF as co-localising are marked with arrows. (H). Venn diagram comparing the number of enriched proteins between 0 (purple) and 24 hour (green) lytic replication TREx cells in the SFPQ (i) and FLAG (ORF11) TMT-LC-MS (ii). Scale bars are 5 µm. All repeats are biological.

Identification of the KSHV ORF11 protein prompted the hypothesis that ORF11 may be the virally-encoded factor driving the formation of these KSHV-induced paraspeckle-like condensates. Like many viral proteins, ORF11 appears to be multifunctional in nature. Previous research has demonstrated that ORF11 is expressed early post reactivation and has been implicated in the formation of specialised ribosomes (39). Therefore, to investigate whether KSHV ORF11 localised to the virus-induced paraspeckle-like structures, FLAG co-immunoprecipitations were initially performed in a TREx-BCBL1-RTA FLAG-ORF11 overexpressing cell line, due to the lack of an available ORF11 antibody, which confirmed an RNA-dependent association between ORF11 and SFPQ (**Fig.3B**). Immunofluorescence studies in the TREx-BCBL1-RTA FLAG-ORF11 cell line also confirmed that ORF11 and SFPQ co-localise in paraspeckle-like condensates at 24 hours post-reactivation, distributed throughout the plane of the cell (**Fig.3C, Fig.S3B**). To further assess ORF11 co-localisation during condensate biogenesis immunostaining was performed during the early stages of lytic replication at 8 and 16 hours. Results confirmed that at 16 hours post lytic induction, ORF11 can be observed co-localising with the immature condensates, as shown through co-localisation with SFPQ (**Fig.3D**). To determine whether KSHV ORF11 was essential for paraspeckle-like condensate formation, a previously characterised CRISPR/Cas9 TREx-BCBL1-RTA ORF11 knockout cell line was utilised(39). As expected, immunofluorescence studies confirmed that SFPQ was not redistributed into puncta or any paraspeckle-like condensates formed in ORF11 knockout cells compared to a scrambled cell line, however early vRTCs were still evident, further supporting a direct role for ORF11 in the biogenesis of the paraspeckle-like condensates (**Fig.3E**).

A classical trait of many paraspeckle proteins is the presence of intrinsically disordered regions (IDRs), which help in the formation of paraspeckles by driving phase separation (40). Notably, structural analysis prediction software (PrDOS and FuzDrop) identified 2 putative IDRs at the N-and C-termini of ORF11, suggesting these domains could be essential for condensate formation (**Fig.3F, Fig.S3C**). Consequently, both termini were deleted in a dual ORF11 truncation mutant, and rescue overexpression studies performed in ORF11-CRISPR cells to discern whether this mutant construct could still maintain the ability to drive paraspeckle condensate formation compared to a wildtype ORF11 construct. Overexpression of the ORF11 truncation mutant failed to rescue condensate formation, in contrast, overexpression of wildtype ORF11 clearly had the ability to drive condensate formation, implicating the ORF11 IDRs in driving paraspeckle-like condensate formation (**Fig.3G**).

Due to the specificity of KSHV ORF11 localising to the virus-induced condensates, affinity pulldowns coupled to TMT-labelled quantitative mass spectrometry were repeated using anti-FLAG or control anti-IgG antibodies in 24 hour reactivated TREx-BCBL1-RTA FLAG-ORF11 overexpressing cells (**Fig.3H, Fig.S3Di-iii**). The interaction profiles were interrogated for presence of essential and core paraspeckle proteins, not surprisingly all early paraspeckle biogenesis factors (5) were present, namely SFPQ, NONO and hnRNP K, indicating ORF11 could play a role in early condensate formation. ORF11 affinity pulldowns also reinforced the interactions observed in the SFPQ affinity pulldowns, highlighting an enrichment of a large number of RNA processing factors not commonly reported to associate with canonical condensates, in particular members of the hnRNP and DEAD/DEAH box helicase families, such as DDX17, DDX21, DDX3X, DDX5, DHX9, hnRNP K and U (**Fig.3H, Fig.S3E)**. Additionally, Venn diagrams (**Fig. 3Ii-ii**) highlight the shift in interaction profile of both SFPQ and ORF11 during lytic replication compared to latent, suggesting a distinct role during lytic replication, with 68 and 62 new interactors for SFPQ and ORF11, respectively at 24 hours. A selection of these proteins highlighted in the TMT-MS, DHX9, hnRNP M, DDX17, DDX21 and hnRNP U, were confirmed to co-localise with the condensates using immunofluorescence studies (**Fig.S3F-J**). Together this data suggests that the KSHV ORF11 protein co-localises to and is required for the formation of virus-induced paraspeckles. Interestingly, we observed a lack of paraspeckle formation in latent ORF11 overexpressing TREx-BCBL1-RTA cells, and localisation of ORF11 to the cellular membrane, suggesting that aadditional viral lytic factors maybe required to localise ORF11 to the nucleus or help to drive paraspeckle biogenesis. Additionally, there are distinct proteomic interactions highlighted in both SFPQ and ORF11 TMT-MS that have not yet been reported in canonical paraspeckles, with some of these factors potentially aiding in biogenesis.

### Paraspeckle-like condensates are essential for KSHV lytic replication

SFPQ is an essential component of canonical paraspeckles and is also exclusively redistributed into the virus-induced paraspeckle-like condensates, therefore loss of SFPQ is predicted to abrogate condensate formation (36). To test this hypothesis, SFPQ stable knockdown cell lines were produced in TREx-BCBL1-RTA cells transduced with targeted shRNAs. Effective SFPQ knockdown reduced SFPQ mRNA levels by 60%, and protein levels by 70%, respectively (**Fig.4A, Fig.S4A-B**). The effect of SFPQ depletion on virus-induced paraspeckle formation was then assessed compared to a scrambled control using NONO-specific antibodies, as the paraspeckle marker. Results confirmed that condensates failed to form in SFPQ-depleted cells compared to control cells (**Fig.4B**). These cell lines were then utilised to assess what effect failure to produce paraspeckle-like condensates had upon KSHV lytic replication. Reactivation of these condensate deficient SFPQ-depleted cells showed a dramatic reduction, of at least 90%, in both the early ORF57 and the late ORF65 mRNA and protein levels (**Fig.4C-E**). To confirm that SFPQ depletion affected KSHV lytic replication, viral load and infectious virion production were also assessed compared to scramble control cells. Viral genomic DNA was measured via qPCR from scrambled and SFPQ-depleted TREx-BCBL1-RTA cells to determine whether viral DNA load was affected. Results showed SFPQ depletion led to a 70% reduction compared to scrambled control (**Fig.4F**). In addition, supernatants of reactivated scrambled and SFPQ-depleted TREx-BCBL1-RTA cells were used to re-infect naïve HEK-293T cells and infectious virion production quantified by qPCR. Cells re-infected with supernatant from SFPQ-depleted cells resulted in a dramatic loss of infectious virions, reduced by 80% compared to controls (**Fig.4G**). Taken together, these results suggest that SFPQ depletion and the resulting failure to form paraspeckle-like condensates significantly impacts KSHV lytic replication and infectious virion production.

**Figure 4:**
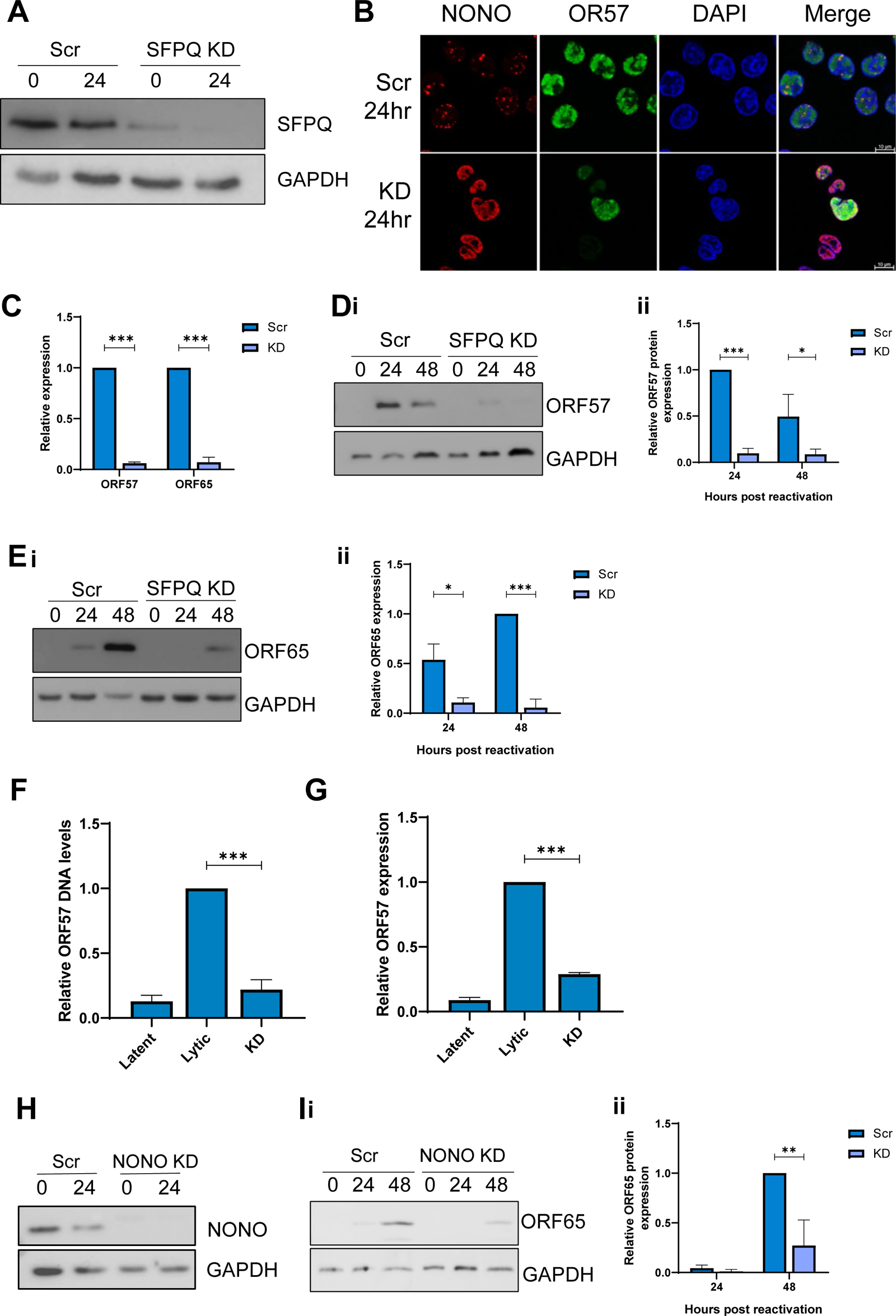
Condensates are essential for viral replication. (A). Representative western blot of SFPQ levels in TREx scr and TREx SFPQ KD cell lines with GAPDH as a loading control (n=3). (B) IF analysis in TREx scr and TREx SFPQ KD cell lines with antibodies against NONO (red) and ORF57 (green). (C) qPCR analysis of ORF57, ORF65 levels in TREx scr and TREx SFPQ KD TREx cells after 24 hours of KSHV lytic replication. GAPDH was used as a housekeeper (n = 3). (D-E) Representative western blots of ORF57 (C) and ORF65 (D) levels in TREx scr and TREx SFPQ KD cell lines with GAPDH as a loading control. Densitometry analysis of (n=3) (Cii and Dii). (F). qPCR analysis of ORF57 DNA levels for viral load at 72 hr post-KSHV induction, with TREx scr and TREx SFPQ KD including uninduced cells as a reactivation control and GAPDH as a housekeeper (n = 3). (G). qPCR analysis of ORF57 in HEK-293T cells for reinfection assay (n = 3). (H). Representative western blot of NONO protein levels in scr and NONO KD TREx cells with GAPDH as a loading control (i). Densitometry analysis of n=3 (ii). (I). Representative western blot of ORF65 protein levels in TREx scr and TREx NONO KD cell lines with GAPDH as a loading control (i). Densitometry analysis of n=3 (ii). Scale bars in B are 10 µm. In C-G, I, data are presented as mean ± SD. *P < 0.05, **P < 0.01 and ***P < 0.001 (unpaired Student’s t-test). All repeats are biological.

SFPQ plays multiple regulatory roles in the nucleus, therefore the essential role of the virus-induced paraspeckle-like condensates was further examined using a stable NONO depleted cell line (**Fig.4H, Fig.S4C-D**), which again showed a significant reduction in viral replication as analysed through ORF65 protein expression, with a 50-70% decrease (**Fig.4I**). IF analysis of NONO KD cells showed the majority of lytic cells failed to form condensates, and any that did form were smaller and malformed compared to the scrambled control (**Fig.S4E**). Furthermore, when *NEAT1* expression was depleted by GapmeRs, condensates once again failed to form in lytic TREx-BCBL1-RTA cells, which resulted in a significant drop in ORF65 protein expression of approximately 75% (**Fig.S4F-H**). Together these findings demonstrate that paraspeckle-like condensate formation is essential for KSHV lytic replication.

### Paraspeckle-like condensates are involved in both viral and cellular RNA processing

The dynamic nature of paraspeckle protein and RNA composition, aligned with the fact that PSPs are generally multifunctional in nature and not exclusively confined to paraspeckles, has complicated the elucidation of paraspeckle function. However, due to the enrichment of RNA helicases and hnRNP members within the virus-induced paraspeckle-like condensates and their adjacent localisation to vRTCs, we hypothesised that the condensates may play a role in viral RNA processing events during KSHV lytic replication. To this end, SFPQ RNA-immunoprecipitations (RIPs) coupled with qPCR were performed in latent and lytic TREx-BCBL1-RTA cells and several viral transcripts were identified as putative SFPQ binders during lytic replication, for example *K8, ORF4,* and *ORF59* (**Fig.5A**). *K8* was the most heavily enriched viral transcript identified in the RIPs, and RNA-FISH was utilised to confirm that *K8* transcripts were localised to the virus-induced paraspeckle-like condensates (**Fig.5B**). Moreover, FLAG-ORF11 RIPs confirmed *K8* association with the condensates (**Fig.5C**). Additionally, SFPQ RIPs were performed in control and ORF11 CRISPR cells to assess association of K8 specificity with condensate formation. Notably the interaction between SFPQ and *NEAT1* was preserved within the CRISPR cells, in contrast, the interaction with *K8* was drastically reduced (**Fig.5D**). This reinforces the observation that the interaction between SFPQ and viral transcripts is dependent on condensate formation.

**Figure 5:**
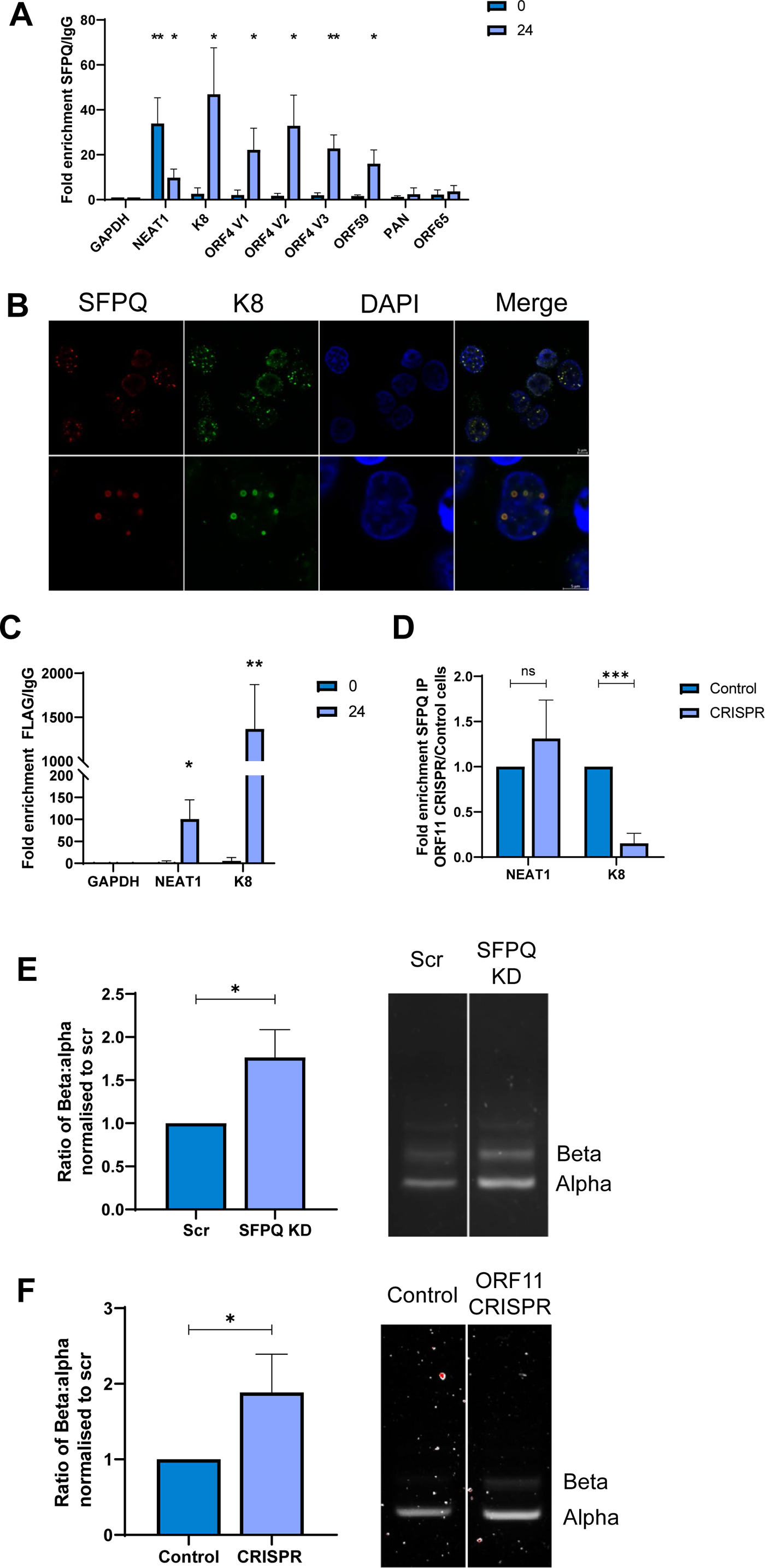
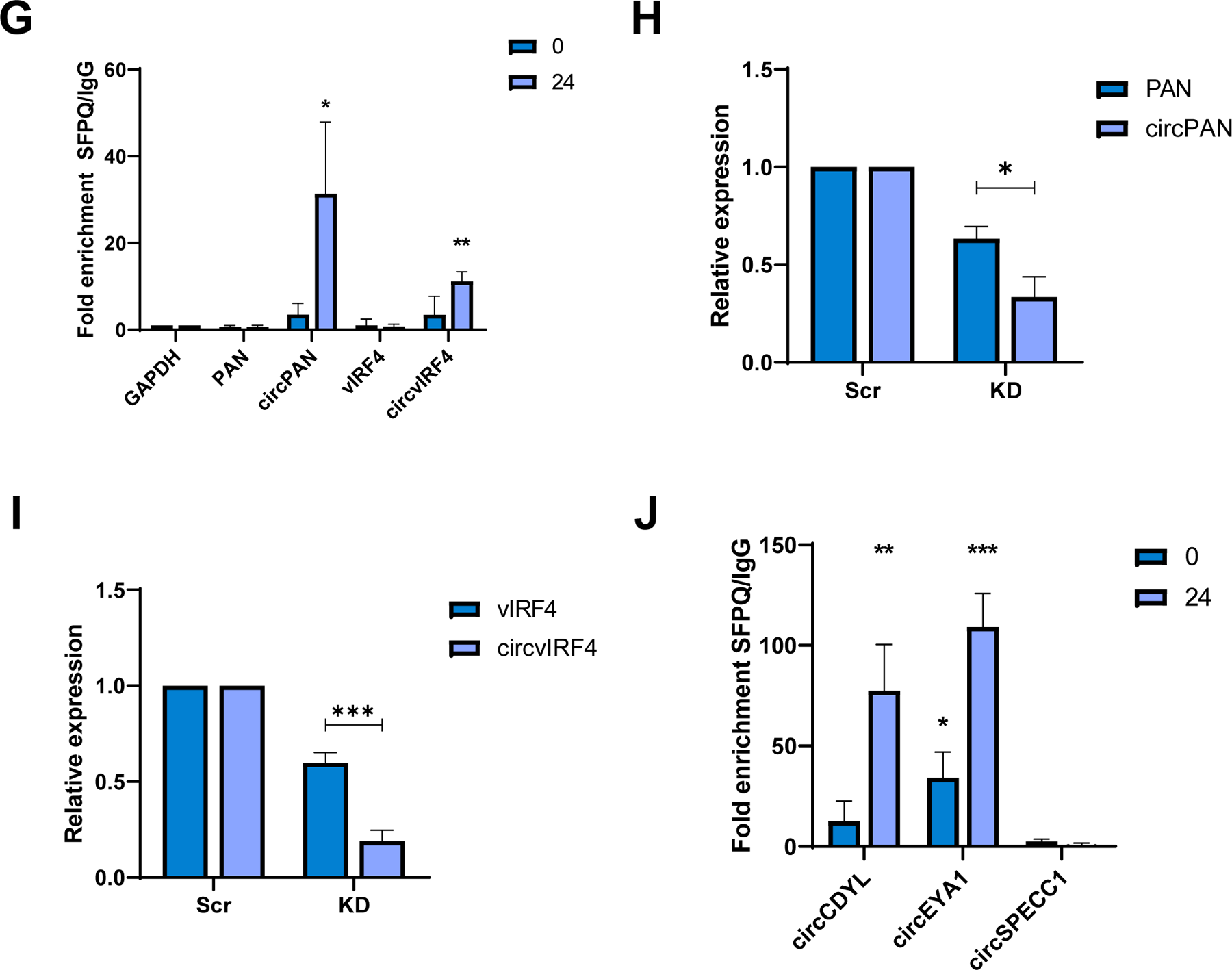
Condensates are implicated in RNA processing. (A) qPCR analysis of SFPQ RIPs in TREx harbouring latent or 24 hours post lytic induction KSHV showing enrichment over IgG normalised to GAPDH (n=3). (B). IF analysis in 24 hour post lytic KSHV induction TREx cells with FISH probes against K8 (green) and antibodies against SFPQ (red). (C). qPCR analysis of FLAG RIPs in TREx FLAG-ORF11 cells at latent and 24 hours post lytic induction showing enrichment over IgG normalised to GAPDH.(n=3). (D). qPCR analysis of SFPQ RIPs in ORF11 CRISPR cells at 24 hours post lytic induction showing enrichment over control cells normalised to NEAT1 (n=3). (E-F) PCR of K8 α, β and γ transcripts in scr and SFPQ KD cells (C) or control and ORF11 CRISPR cells (D). Ratio of α to β was normalised to scr control. Densitometry analysis performed on n=3. (G). qPCR analysis of SFPQ RIPs in TREx cells at latent and 24 hours post lytic induction showing enrichment over IgG normalised to GAPDH (n=3). (H). qPCR analysis of Pan and circPAN levels in scr and SFPQ KD TREx cells at 24 hours post lytic induction. GAPDH was used as a housekeeper (n = 3). (I). qPCR analysis of IRF4 and circIRF4 levels in scr and SFPQ KD TREx cells at 24 hours post lytic induction. GAPDH was used as a housekeeper (n = 3). (J). qPCR analysis of SFPQ RIPs in TREx cells at latent and 24 hours post lytic induction showing enrichment over IgG normalised to GAPDH(n=3). Scale bars are 5 µm. In A, C-J data are presented as mean ± SD. *P < 0.05, **P < 0.01 and ***P < 0.001 (unpaired Student’s t-test). All repeats are biological.

The *K8* transcript undergoes alternative splicing to produce three spliced variants, termed α, β and γ. It has been previously demonstrated that the majority of the pre-mRNA γ transcript is processed into the functional α transcript and a small proportion of transcripts retain an intron, becoming the β variant which is not translated (41, 42). Although the exact role of these alternative spliced variants is unknown, maintaining this ratio is critical for KSHV lytic replication, with the β truncated protein potentially acting as an antagonist for the primary K8 protein (41–43). To elucidate whether virus-induced paraspeckle-like condensates have a role in orchestrating *K8* processing, a rudimentary PCR assay was performed to analyse the effect condensate formation had on the production of alternative *K8* spliced variants. Here *K8* splicing was assessed in the non-condensate forming SFPQ-depleted and ORF11 knockout cell lines compared to scrambled controls. Notably, in both cell lines which fail to form paraspeckle-like condensates, enhanced levels of the non-translated *K8* β transcript were observed compared to baseline α levels (**Fig.5E-F**). This implicates loss of condensate formation in the dysregulation of viral RNA processing.

Recently SFPQ has also been postulated to have a role in circRNA biogenesis (12). Therefore, aligned with the recent observation that KSHV encodes its own circRNAs (kcircRNAs) and also manipulates host circRNA levels during lytic replication (44, 45), a potential association in the biogenesis and localisation of circRNAs to condensates was investigated. Although many kcircRNAs have been identified through sequencing, two of the most abundant are kcircPAN and kcircvIRF4 (44, 46, 47). Notably, SFPQ RIP coupled with qPCR showed a clear association with both kcircRNAs (**Fig.5G**) and interestingly their levels were disproportionately downregulated over their linear transcripts in SFPQ-depleted cells (**Fig.5H-I**). Moreover, to determine whether condensates could also act as processing hubs for cellular circRNAs co-opted by KSHV, SFPQ RIPs were performed to identify interacting host circRNAs and whether their levels were affected by SFPQ depletion. Both circCDYL and circEYA1 were identified as enriched within SFPQ RIPs coupled with qPCR (**Fig.5J**), confirming recent results (12). To determine whether these SFPQ-interacting cellular circRNAs had any specific role in virus replication, qPCR analysis was used to assess any changes in levels during replication and results showed that both circCDYL and circEYA1 levels were increased during lytic replication, independently of their parental linear transcripts (**Fig.S5A-B**). Importantly, shRNA-mediated depletion of either circEYA1 or circCDYL resulted in a significant loss of the late viral ORF65 protein, thus highlighting the importance of these circRNAs to KSHV lytic replication (**Fig. S5E-G**). Finally, similar to the KSHV-encoded circRNAs this upregulation was significantly impacted by SFPQ depletion (**Fig.S5C-D**). Together these results suggest that paraspeckle-like condensate formation and SFPQ sequestration is important for both viral and host cell circRNA biogenesis during lytic replication.

### Large paraspeckle-like condensates are formed in gammaherpesvirus infection

To examine whether the novel virus-induced paraspeckle-like condensates identified during KSHV lytic infection are formed during other herpesvirus infections, immunofluorescence studies were performed in human foreskin fibroblast (HFF) cells infected with either the alphaherpesvirus Herpes simplex virus type 1 (HSV-1) or the betaherpesvirus human cytomegalovirus (HCMV). Immunostaining with SFPQ-specific antibodies was performed at 0, 4 and 8 hours post-infection for HSV-1 and 0, 48 and 72 hr post-infection for HCMV, respectively. However, in contrast to KSHV-infected cells, no redistribution of SFPQ was observed into large puncta during either infection (**Fig.6A-B**). Therefore, to determine whether paraspeckle-like condensates are unique to KSHV, or occurred in other gamma-herpesvirus infections, immunofluorescence studies were repeated in Epstein Barr Virus (EBV)-infected cells: 293-rJJ-L3s, which harbours a BAC EBV clone (the type 2 EBV Jijoye strain), which were reactivated via transfection of the EBV lytic proteins ZTA, RTA and BALF4 (48, 49). Similar to KSHV, SFPQ re-localised from a diffuse nuclear staining observed during latency to form puncta that clustered around EBV replication compartments, indicated by Ea-D staining, at 24 and 48 hours post-lytic induction (**Fig. 6C**). These results suggest that SFPQ containing puncta are only formed during gamma herpesvirus lytic replication cycles.

**Figure 6:**
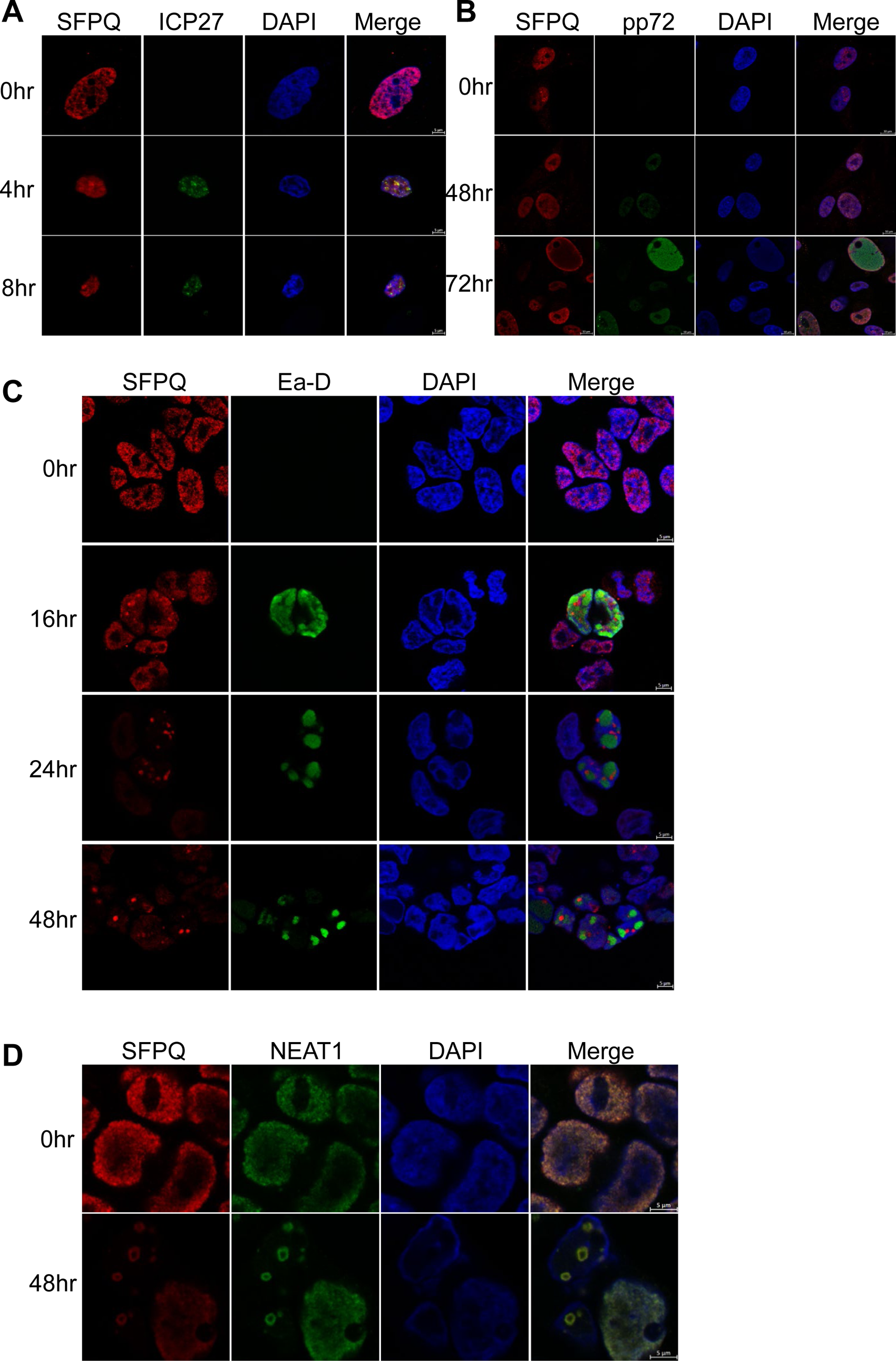
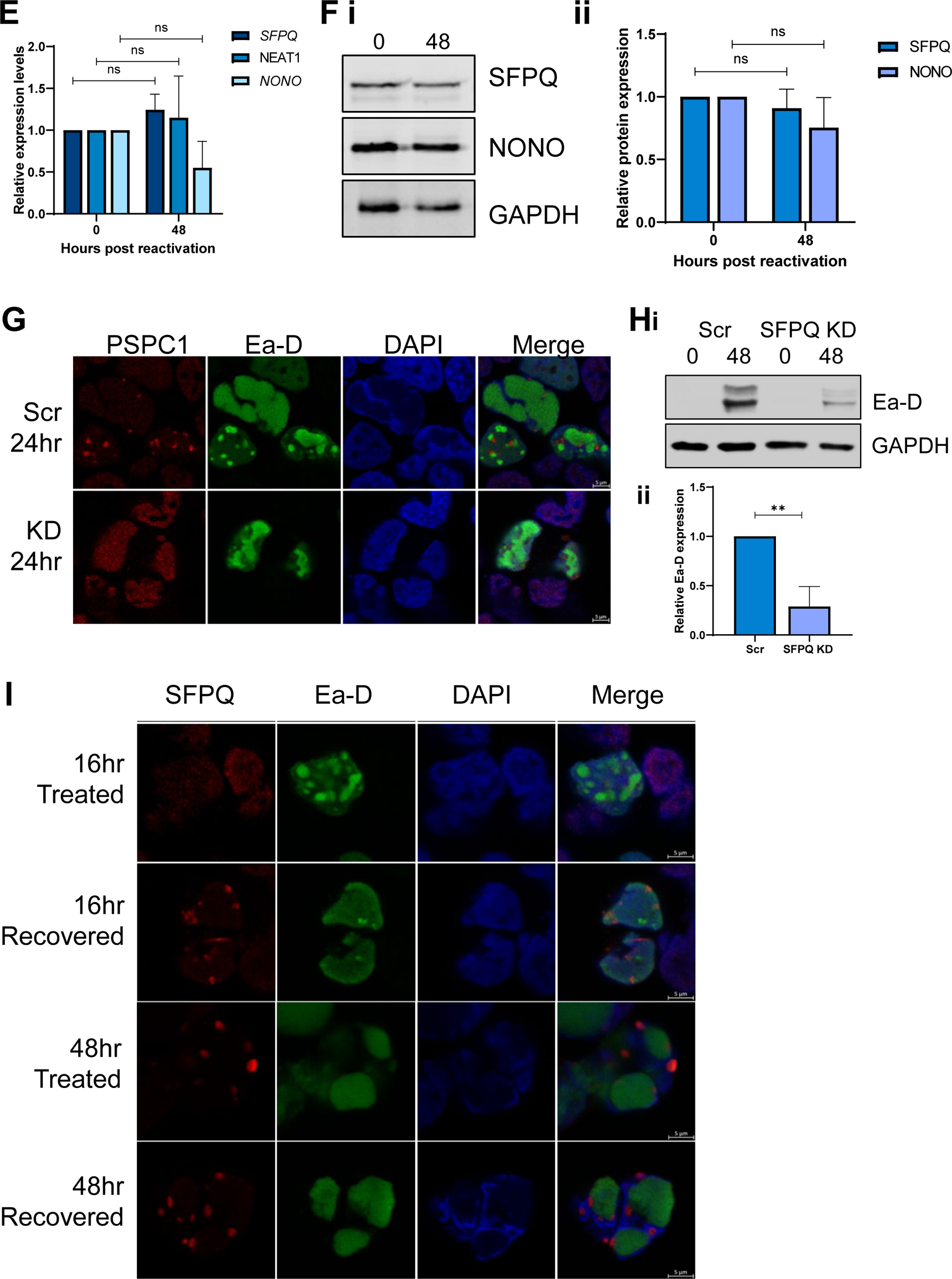
Large paraspeckle-like condensates are only formed in gamma-herpesvirus lytic replication. (A) IF of HFF cells infected with HSV-1 at 0, 4 and 8 hours, stained for ICP27 and SFPQ. IF of HFF cells infected with HCMV at 0, 48 and 72 hours, stained for SFPQ and pp72. (C): IF of 293-rJJ-L3s cells at 0, 16, 24 and 48 hours post-reactivation with antibodies against SFPQ (red) and EA-D (green). (D). IF of 293-rJJ-L3s cells at 48 hours reactivation, stained for SFPQ (red) and NEAT1 (green). (E). qPCR of SFPQ, NONO and NEAT1 levels in rJJ-L3-#1 cells at 0 and 48 hours. GAPDH was used as a housekeeper, n=3. (F). Representative western blot SFPQ and NONO protein levels in 293-rJJ-L3s cells at 0 and 48 hours with GAPDH as a loading control (i). Densitometry analysis of n=3 (ii). (G). IF analysis in scr and SFPQ KD rJJ-L3-#1 cells with antibodies against PSPC1 (red) and Ea-D (green). (H). Representative western blot of Ea-D protein levels in scr and SFPQ KD 293-rJJ-L3s cells with GAPDH as a loading control (i). Densitometry analysis of n=3 (ii). (I). IF of 293-rJJ-L3s cells at 16 and 48 hours. Cells were either treated with 1,6 HD or treated and then allowed to recover for 30 minutes prior to fixing. Scale bars are 5 µm in all IF except B (10 µm). In E-F, H data are presented as mean ± SD. *P < 0.05, **P < 0.01 and ***P < 0.001 (unpaired Student’s t-test). All repeats are biological.

Similar to KSHV, the SFPQ puncta formed during EBV lytic replication were large non-canonical ring-like structures and RNA FISH confirmed the co-localisation of *NEAT1*, again suggesting the formation of virus-induced paraspeckle-like puncta (**Fig.6D**). Moreover, both SFPQ and NONO mRNA and protein expression, as well as *NEAT1* RNA levels, were stable during EBV lytic replication suggesting an alternative biogenesis pathway to canonical paraspeckles, analogous to KSHV infection (**Fig.6E-F**). To determine whether formation of these puncta was also required for EBV lytic replication, SFPQ was depleted in the 293-rJJ-L3s cell line and immunofluorescence studies using PSPC1 confirmed that SFPQ was required for paraspeckle-like puncta formation during EBV lytic replication (**Fig.6G**). Moreover, SFPQ depletion resulted in a dysregulation of the early EBV lytic cycle, as shown by a 70% reduction of Ea-D protein expression (**Fig.6H**). Finally, 1,6-hexanediol treatment experiments showed a maturation process from a liquid-like state during early EBV lytic replication (16 hours) to a more gel-like state during the later stages of infection (48 hours) (**Fig.6I**). Together, these data suggest that large paraspeckle-like condensates are potentially a pan-gammaherpesvirus phenomenon that occur during the lytic replication phase.

### Formation of virus-induced paraspeckles-like condensates impact on genomic instability

The finding that paraspeckle-like condensate formation was only observed in gammaherpesvirus infection is particularly intriguing and led to further studies exploring a potential link between paraspeckle-like condensate formation and gammaherpesvirus-mediated tumourgenesis. Recent observations have highlighted that both KSHV and EBV lytic replication cycles contribute to an increase in genome instability, although the mechanisms driving virus-mediated tumourgenesis are not fully elucidated (27, 50, 51). Notably, a potential role of SFPQ has also been postulated in the DNA damage response (DDR) (52–54), which suggests an intriguing link between the increase in genomic instability and the formation of the virus-induced paraspeckle-like condensates, due to increased sequestration and association of PSPs, resulting in their depletion from the nucleoplasm. This is particularly evident with SFPQ which shows loss of the diffuse staining throughout the nucleoplasm upon the formation of virus-induced paraspeckle-like condensates.

A key marker of DNA damage is lagging chromosomes during mitosis, where chromosomes fail to separate into daughter cells. Upon investigation of the absence or presence of chromosomal anomalies in scrambled vs SFPQ depleted TREX-BCBL1-RTA or HEK293T cells, the percentage of cells with lagging chromosomes increased from 0% to 30%, and from 0% to 50%, respectively upon depletion of SFPQ (**Fig.7A**). This implicates SPFQ as a factor in the DDR, therefore depletion of SFPQ could lead to increases in DNA damage.

**Figure 7:**
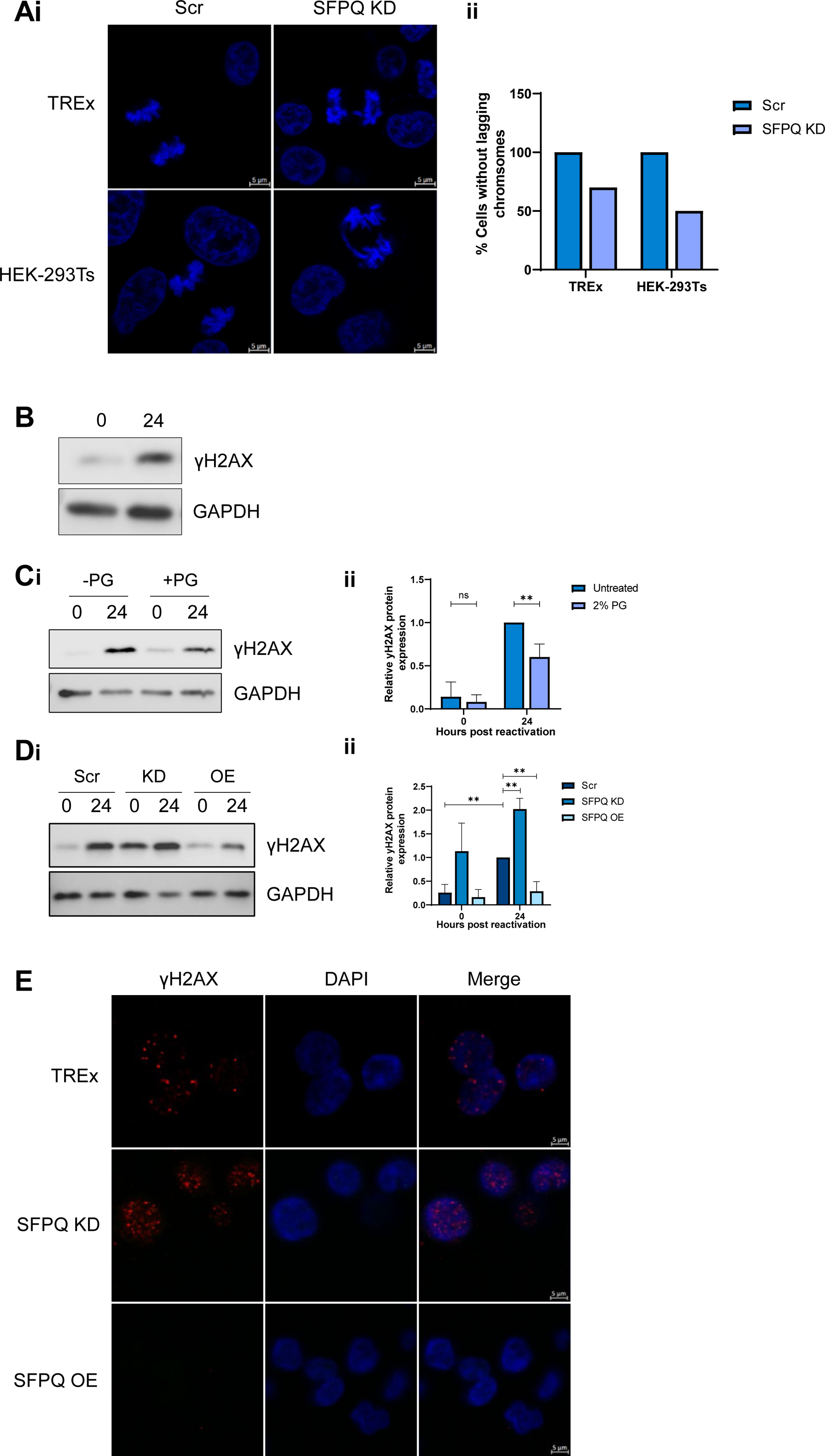
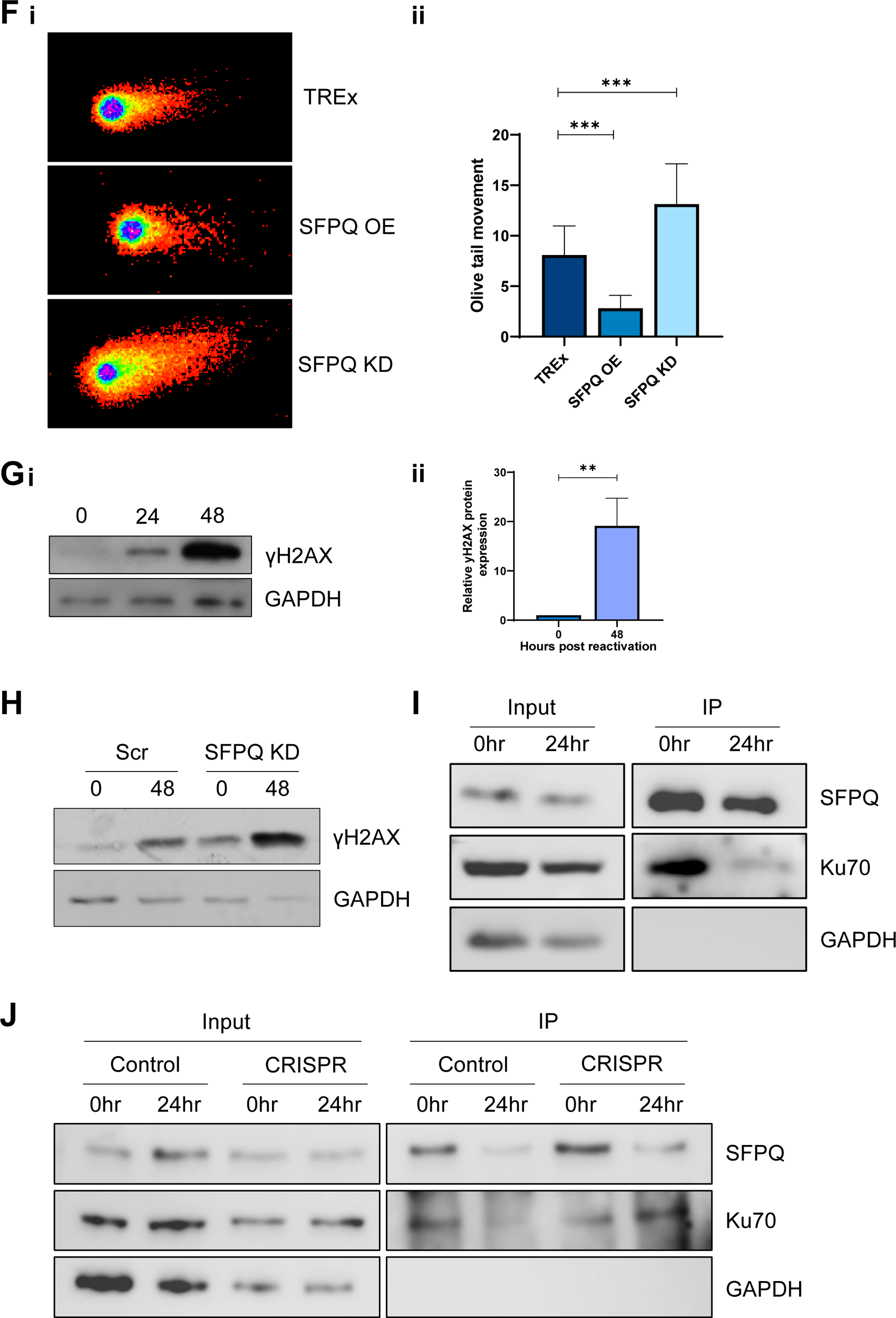
Formation of condensates disrupts the DDR. (A). IF of scr and SFPQ KD latent TREx and HEK-293Ts (i). 20 cells undergoing active mitosis were counted for each cell type to identify the percentage of lagging chromosomes (ii). (B). Representative western blot for levels of γH2AX in TREx cells at latent and 24 hours post-lytic induction. GAPDH was used as a loading control (n=3). (C). Representative western blot for levels of γH2AX in TREx cells at latent and 24 hours post lytic induction (i). Cells were treated with propylene glycol (PG) between 16 and 24 hours post-reactivation. GAPDH was used as a loading control. Densitometry analysis was performed on n=3 (ii). (D). Representative western blot of γH2AX in scr, SFPQ KD and SFPQ OE TREx at latent and 24 hours post lytic induction (i). GAPDH was used as a loading control, densitometry analysis was performed on n=3 (ii). (E). IF of γH2AX (red) in latent scr, SFPQ KD and SFPQ OE TREx. (F). Representative neutral comet assays for scr, SFPQ O/E and SFPQ KD TREx during latency (i). Olive tail movement was calculated using Comet Score 2.0 (n=3), with 50 cells counted per sample per biological repeat. (G). Representative western blot for levels of γH2AX in 293-rJJ-L3s cells at latent and 24 and 48 hours post-lytic induction (i). GAPDH was used as a loading control, densitometry analysis for latent and 48 hours post-lytic induction was performed on n=3 (ii). (H). Representative western blot for levels of γH2AX in scr and SFPQ KD 293-rJJ-L3s cells at latent and 48 hours post-lytic induction. (I). Co-IP of SFPQ in TREx cells at latent and 24 hours post-lytic induction probed with antibodies against Ku70 and GAPDH (n=3). (J). Co-IP of SFPQ probed with antibodies against Ku70 and GAPDH (n=3) in control and CRISPR ORF11 at latent and 24 hours post-lytic induction. Scale bars are 5 µm. In C-D, F-G data are presented as mean ± SD. *P < 0.05, **P < 0.01 and ***P < 0.001 (unpaired Student’s t-test). All repeats are biological.

As previously described, KSHV lytic replication leads to an increase in markers of DNA damage, as measured by immunoblotting using γH2AX-specific antibodies, a marker of double stranded DNA breaks (**Fig.7B**). To determine whether this increase in DNA damage is linked to SFPQ sequestration, condensate formation was disrupted during KSHV lytic replication using propylene glycol (PG). This has a similar mechanism to 1,6-hexanediol, allowing disruption of condensates in a liquid-like state, but can be used for longer time periods after hyperosmotic adaptation (38). TREx-BCBL1-RTA cells were incubated with PG between 16-24 hours post-lytic induction, leading to an environment in which viral replication could partly occur, however condensate formation was abrogated (**Fig.S6A**). Under these conditions, SFPQ remained in the nucleoplasm and not recruited to virus-induced paraspeckle-like condensates. To determine whether there is a link between condensate formation and genomic instability, DNA damage was then compared in KSHV latent and lytically replicating cells, in the absence and presence of PG. 23]. Interestingly, however, in the presence of PG, a significant reduction in γH2AX levels were observed compared to control reactivated cells. This suggests that reduced SFPQ sequestration into the virus-induced condensates may prevent virus-induced genomic instability (**Fig.7C**). To confirm this observation, DNA damage was compared in the KSHV ORF11 knockout CRISPR cells, where condensates are disrupted but early lytic viral gene expression is still evident. Results show a similar phenotype to PG treatment with reduced γH2AX levels in ORF11 knockdown cells compared to control (**Fig.S6B**).

To confirm that the increase in DNA damage observed due to virus-induced paraspeckle-like condensate formation was specifically due to SFPQ sequestration, changes in γH2AX abundance were compared in latent and reactivated TREx-BCBL1-RTA cells either depleted for SFPQ or upon SFPQ overexpression. As expected, depletion of SFPQ resulted in an increase in γH2AX, whereas overexpression of SFPQ, which provides sufficient SFPQ to function in the nucleoplasm as well as being sequestered in virus-induced condensates, led to reduced γH2AX levels regardless of whether KSHV-infected cells remained latent or underwent lytic replication (**Fig.7D-E**). Similarly, this increase in DNA damage markers was also observed during EBV lytic replication, with levels of γH2AX again increasing during lytic replication and inversely correlating with SFPQ levels (**Fig.7G-H, Fig.S6C**).

Neutral comet assays were then used to quantify double stranded breaks in scrambled latent control cells compared to TREx-BCBL1-RTA cells either depleted for SFPQ or upon SFPQ overexpression. A similar phenotype was observed, where SFPQ overexpressing cells had the lowest mean olive tail movement (OTM) of 2.8 equating to the least DNA damage, followed by scrambled control cells (8.1), whilst SFPQ KD cells had the greatest mean OTM of 13.1 and therefore the greatest number of dsDNA breaks (**Fig.7F**). Together, these results validate the hypothesis that gammaherpesvirus-induced condensate and SFPQ sequestration contribute to tumourigenesis.

Finally, the potential role of SFPQ sequestration into virus-induced paraspeckle-like condensates and increased genomic instability was investigated. It has been previously observed that SFPQ can interact with the Ku70/80 dimer and act as a scaffold for the pre-ligation complex, improving effectiveness of the DDR machinery, whilst Ku70 was also enriched within the SFPQ TMT-MS (36, 52, 53). Therefore, the interaction between SFPQ and Ku70 was assessed in KSHV latent and lytic TREx-BCBL1-RTA cells. Co-immunoprecipitation assays confirmed an interaction between SFPQ and Ku70 in latent cells, however, this interaction was significantly decreased during KSHV lytic replication (**Fig.7I**). Conversely, when virus-induced paraspeckle-like condensates do not form (using the ORF11 CRISPR cell line) the interaction between SFPQ and Ku70 is increased compared to the scramble control (**Fig.7J**). Together, these data suggest that formation of virus-induced paraspeckle-like condensates leads to an increase in dsDNA breaks due to SFPQ sequestration, which in turn may contribute to gammaherpesvirus oncogenesis.

In summary, these findings identify SFPQ and the KSHV-encoded protein ORF11 as key drivers in the formation of novel virally-induced paraspeckle-like condensates which act as hubs for both viral and cellular RNA processing during KSHV lytic replication. Formation of these novel condensates also promote genomic instability during lytic replication due to SFPQ sequestration, by preventing its interaction with Ku70 and potentially affecting recruitment of the pre-ligation complex in the DDR. This highlights a novel mechanism which increases DNA damage during gammaherpesvirus lytic replication which may be associated with viral-mediated tumourigenesis.

## Discussion

The dynamic nature of biomolecular condensates allows fast changes to cellular stimuli, enabling a fine-tuned cellular response to a range of conditions in a compartmentalised manner. However, they are also frequently dysregulated in disease states such as cancer, Alzheimer’s disease and fronto-temporal dementia (55–58). Wider research has demonstrated viruses are able to either induce or alternatively disrupt condensate formation to enhance their replicative cycles, such as inhibition of stress granules or formation of viral factories (38, 59). This study has identified a novel gammaherpesvirus-induced paraspeckle-like condensate, which acts as an essential hub for both viral and cellular RNA processing during infection. Critically, formation of these structures are essential for successful viral lytic replication, whilst simultaneously increasing DNA damage through sequestration of PSP proteins, thus potentially contributing to gammaherpesvirus-mediated tumourigenesis.

The establishment of the virus-induced paraspeckle-like condensates occurs through liquid-liquid phase separation, initially forming a dynamic, liquid-like body that matures into a gel-like state over the first 24 hours of viral lytic replication. Importantly, although they utilise many canonical paraspeckle components including *NEAT1*, SFPQ and NONO, there are several key differences, marking them as paraspeckle-like. The first major difference is highlighted by high-resolution microscopy which showed virus-induced paraspeckles-like condensates are up to 10x larger than canonical paraspeckles and, importantly, their shape remains highly spherical, which contrasts with the elongation model previously observed in canonical paraspeckles. Secondly, canonical paraspeckle formation occurs co-transcriptionally with *NEAT1* expression correlating with number or size. *NEAT1* containing RNPs then form through binding of SFPQ and NONO, with multiple RNPs then being joined through FUS to form a mature paraspeckle. Surprisingly however, qPCR analysis confirmed *NEAT1* levels remain stable throughout KSHV viral replication despite the increase in paraspeckle-like number and size. Moreover, a combination of TMT-MS analysis, IF and Co-IP studies highlighted several of the core PS proteins are missing, most notably RBM14, and FUS, with IF confirmed FUS is re-localised to the vRTCs. Notably, loss of core PS proteins is known to inhibit canonical paraspeckle formation (60, 61), conversely formation of virus-induced paraspeckle-like condensates appear to be independent of these factors, implicating a distinct architecture and biogenesis pathway that may utilise different factors or mechanisms.

Alternative biogenesis of the virus-induced paraspeckle-like condensates is likely to be mediated through the KSHV-encoded protein ORF11, which co-localises with SFPQ throughout the entirety of virus-induced paraspeckles formation and lifetime. Cells lacking ORF11 fail to form virus-induced paraspeckles and importantly, like many proteins that drive phase separation, ORF11 has low complexity terminal ends, with truncation of these ends inhibiting virus-induced paraspeckle formation. Further work aims to elucidate the exact mechanism ORF11 plays in virus-induced paraspeckle formation, whether it is a direct FUS replacement or plays an alternative role. Whilst similar paraspeckle-like condensates form in EBV, the virus does not contain a direct ORF11 homologue and the closest paralogue to ORF11 in all human herpesviruses, the EBV protein LF2, has only ∼28% protein blast alignment sequence similarity. It will be of interest to determine whether LF2 plays a similar role to KSHV ORF11 or whether an alternative viral factor is involved in EBV-induced paraspeckle-like formation.

It is clear virus-induced paraspeckle-like condensates are essential for viral replication, with depletion of SFPQ, NONO or *NEAT1* leading to significant reductions in the levels of KSHV lytically expressed RNAs and proteins. This reduction is likely due to paraspeckle-like condensates playing a critical role in the processing of viral transcripts, as exemplified for K8 processing, essential for viral replication. It is also interesting to note that the levels of both virally-encoded and cellular circRNAs are downregulated upon paraspeckle-like condensation disruption. Although the exact role is still to be elucidated, the enrichment of RNA helicases and hnRNPs in quantitative MS combined with enrichment of these circRNAs in SFPQ-RIP analysis highlights a possible of sequestration of these RNA processing factors in circRNA biogenesis. This is supported by recent findings showing that the RNA helicase DDX5 in combination with the m6A reader YTHDC1, both of which are enriched within the TMT LC-MS/MS, have been found to specifically enrich the backsplicing of specific circRNAs (62). Furthermore, several of these circRNAs have long flanking introns and have been shown implicated as regulated by SFPQ (12). Similarly, hnRNP M has also been shown to positively regulate the biogenesis of circRNAs with long flanking introns (63).

One future direction of this work is to determine the targeting process into virus-induced paraspeckle-like structures, as only certain viral and cellular transcripts are associated in the condensates. Many of these viral transcripts are intron-containing whilst there is also an enrichment of circRNAs with long flanking introns. Previous research has highlighted RNAs enriched in paraspeckles or processed by SFPQ often have these features (6, 64), thus this may be a key determining factor. Additionally, the nuclear m^6^A reader YTHDC1, which is critical for directing methylated transcripts to distinct biological fates, alongside its cofactors hnRNP A2B1 and hnRNP C was identified as a novel SFPQ interactor within the TMT LC-MS/MS. Thus, m^6^A methylation may play a role in determining which transcripts are associated with the paraspeckle-like structures as many of the viral and cellular transcripts identified are heavily methylated (65–67). The proximity of this complex suggests it could be orchestrating the translocation of viral transcripts from vRTCs directly to the paraspeckle-like condensates via targeted methylation.

Due to the specificity of condensate formation for oncogenic gammaherpesviruses and the observed increase in chromosomal abnormality upon SFPQ depletion, we further explored a potential link between virus-induced paraspeckle formation and genomic instability. An increase in genomic instability during both KSHV and EBV lytic replication has been previously reported, however mechanisms driving DNA damage are yet to be fully elucidated. Both latent and lytic life cycles are essential for virus-mediated tumourgenesis, and specifically regarding lytic replication, it is proposed that over time, the continuous reactivation in infected individuals either undergoing the full lytic cycle or abortive lytic replication leads to increased DNA damage, thus increasing the risk of cancer development. For example, previously, it has been shown that the sequestration of hTREX components to viral mRNAs drives R-loop formation as a contributing factor whilst several DDR proteins including RPA32 and MRE11 have been shown to localise to vRTCs (24, 68). Herein, we identify a novel mechanism which contributes to genomic instability in both KSHV and EBV lytic replications, namely the sequestration of SFPQ into virus-induced paraspeckle-like condensates resulting in a concomitant increase in DNA damage. Therefore, combined with enhanced R-loop formation associated with virus-mediated hTREX sequestration, the proposed sequestration of SFPQ into the paraspeckle-like condensate would lead to a ‘dual hit’ on DNA damage, with the DDR response potentially also being impaired. SFPQ expression inversely correlates with accumulation of DNA damage, specifically DSBs. SFPQ is a known interactor with the Ku proteins, key players in NHEJ, which is the main repair pathway during DSB. During lytic viral replication, this interaction is reduced, likely resulting in reduced efficacy of downstream parts of the repair pathway.

To summarise, gammaherpesviruses manipulate core paraspeckle components to form novel condensates, which act as critical hubs for successful viral replication through RNA processing roles. Specifically, KSHV utilises the virally-encoded protein ORF11 as a driver for condensate initiation and formation. Interestingly, the sequestration of SFPQ into the condensates results in increased DNA damage, which may contribute to gammaherpesvirus oncogenesis. Finally, targeting these condensates therapeutically may both target viral lytic replication and additionally reduce the risk of cancer development.

## Methods

### Cell culture

TREx-BCBL1-RTA cells, a gift from Professor JU Jung (University of Southern California), a B-cell lymphoma cell line latently infected KSHV engineered to contain a doxycycline-inducible myc-RTA were cultured in RPMI 1640 with glutamine (Gibco), supplemented with 1% P/S (Gibco), 10% FBS (Gibco) and 100 µg/ml hygromycin B (ThermoFisher). KD cell lines were additionally cultured with 3 µg/ml puromycin (Gibco). ORF11-CRISPR cells have been previously described (Murphy et al., 2023). HEK-293T cells were purchased from the ATCC and cultured in DMEM (Lonza) and supplemented with 10% FBS and 1% P/S. HEK-293T-rKSHV.219 were kindly provided by Dr Jeffery Vieira (University of Washington) and were cultured in DMEM (Lonza), supplemented with 10% FBS, 1% P/S and 3 μg/ml puromycin (Gibco). Human foreskin fibroblasts (HFFs) were a gift from J. Sinclair (University of Cambridge) and cultured in DMEM with 10% FBS and 1% P/S. 293-rJJ-L3 cells are 293 clone #19 carrying a GFP-negative rJJ-L3 Jijoye EBV BAC were cultured in RPMI 1640 with glutamine (Gibco), supplemented with 1% P/S (Gibco), 10% FBS (Gibco) and 100 µg/ml hygromycin B (ThermoFisher). HCMV (Merlin) and HSV-1 virus (SC16) stocks were provided by J. Sinclair and S. Efstathiou (University of Cambridge). All cell lines tested negative for mycoplasma. All cell lines were cultured at 37°C at 5% CO_2_.

Virus lytic replication in TREx-BCBL1-RTA cells was induced via addition of 2 µg/ml doxycycline hyclate (Sigma-Aldrich). HEK-293T-rKSHV.219 cells were induced via addition of 20 ng/ml TPA and 3 mM sodium butyrate. BCBLs were induced via addition of 1.5 mM sodium butyrate. EBV cell lines were trypsinised and induced by addition of even amounts of pBZLF1, pIE-Rta and pBALF4 plasmids using Lipofectamine 2000 (ThermoFisher) and Opti-MEM (ThermoFisher) during seeding.

100nM NEAT1 GapmeRs (antisense complementary oligonucleotides that induce target degradation through RNAse H recruitment) (Qiagen) were added to cells for 24 hours prior to reactivation. For 1,6-hexanediol treatment, cells were subjected to one of three conditions: protection, treatment or recovery. For all conditions a 3% v/v treatment of 1,6-hexanediol from a 1 M stock was added to cells for 30 seconds. Protected cells were treated with 4% (v/v) paraformaldehyde for 5 minutes prior to 1,6-hexanediol treatment, whilst recovered cells had media replaced after the 1,6-hexanediol treatment followed by a 30 minute incubation at 37℃. All conditions were washed with PBS followed by fixation. For propylene glycol treatment, cells were passaged in hyperosmotic conditions before addition of 4% of propylene glycol for 8 hours.

### Plasmid and antibodies

Antibodies used are listed below: GAPDH (Proteintech 60004-1-Ig, WB 1/5000), ORF57 (Santa Cruz sc-135747, WB 1/1000, IF 1/100), ORF65 (CRB crb2005224, 1/100), SFPQ (Proteintech 15585-1-AP, WB 1/500, IF 1/50, RIP 1/50, IP 1/50), NONO (Proteintech 11058-1-AP, WB 1/1000, IF 1/100), PSPC1 (Proteintech 16714-1-AP, WB 1/3000, IF 1/50) FUS (Proteintech 11570-1-AP, WB 1/5000, IF 1/100), FLAG (Sigma F7425, WB 1/500), GFP (Proteintech 66002-1-Ig, WB 1/5000), Ku70 (Proteintech 10723-1-AP, WB 1/2000), γH2AX (CST, WB 1/1000, IF 1/100), SRSF2 (Novus Bio NB100-1774SS, IF 1/250), RNA pol II (Sigma-Aldrich 05-623, IF 1/50), SFPQ (Proteintech 67129-1-Ig, IF 1/400, IP 1/400), FLAG (Sigma F1804, IF 1/50), hnRNP U (Proteintech 14599-1-AP, IF 1/20), hnRNP M (Proteintech 26897-1-AP, IF 1/50), DDX17 (Proteintech 19910-1-AP, IF 1/10), DDX21 (Proteintech 66925-1-Ig, IF 1/50), DHX9 (Proteintech 67153-1-Ig, IF 1/50), BUD23 (Proteintech PA521698, IF 1/50), PNO1 (Proteintech 21059-1-AP, IF 1/50).

pVSV.G and psPAX2 were a gift from Dr Edwin Chen (University of Leeds). PLKO.1 TRC cloning vector was bought from Addgene (gift from David Root; Addgene plasmid #10878). FLAG-ORF11 OE and ORF11 CRISPR plasmids have been described previously (39). ORF11 truncation mutant was cloned via PCR amplification of truncated form of ORF11 from TREx-BCBL1-RTA cell cDNA and cloned used NEBuilder HIFI DNA assembly kit (NEB) into pLenti-CMV-GFP Zeo plasmid (purchased from addgene #17449).

GFP-SFPQ OE plasmid was generated via PCR amplification of SFPQ from TREx-BCBL1-RTA cell cDNA and cloned using NEBuilder HIFI DNA assembly kit (NEB) into pLenti-CMV-GFP-puro plasmid (purchased from addgene #17448). pBZLF1, pIE-Rta and pBAL4 were provided by Dr Rob White (ICL).

### Lentivirus-based shRNA KD

Lentiviruses were generated by transfection of HEK-293Ts, as previously described (45). Per single 6-well, 4 µl Lipofectamine 2000 (ThermoFisher) was used in combination with 1.2 µg pLKO.1 plasmid expressing the shRNA, 0.65 µg psPAX2 and 0.65 µg pVSV.G. 2 days post-transfection, supernatants were collected and filtered using a 0.45 µm filter and used for transductions on the target cells, in the presence of 8 µg/ml polybrene (Merck Millipore). 3 µg/ml puromycin (Gibco) was added 48 hours post-transduction, before KD analysis via qPCR and WB if appropriate.

### RNA extraction and qPCR

Total RNA was extracted using Monarch Total RNA Miniprep kit (NEB) as per instructions. 1 µg RNA was reverse transcribed using LunaScript RT SuperMix Kit (NEB). qPCR was performed utilising appropriate primers, cDNA and GoTaq qPCR MasterMix (Promega) on Rotorgene Q (Qiagen) and analysed via ΔΔ CT against a housekeeping gene as previously described (69). Primers are listed in a supplementary table.

### Immunoblotting

Cell lysates were separated using 8-15% polyacrylamide gels and transferred to Amersham Nitrocellulose Membranes (GE healthcare) via Trans-blot Turbo Transfer system (Bio-Rad). Membranes were blocked in 5% milk in 1xTBS-T or 5% BSA in 1xTBS-T dependent on primary antibody. Membranes were probed with appropriate primary antibodies before secondary IgG HRP conjugated antibodies (Dako Agilent). Proteins were detected with ECL Western Blotting Substrate (Promega) or SuperSignal™ West Femto Maximum Sensitivity Substrate (ThermoFisher) before visualisation with G box (Syngene).

### Immunofluorescence and Fluorescence *in situ* hybridisation (FISH)

Cells were seeded onto coverslips. For suspension cells, coverslips were pre-treated with poly-L-lysine and cells allowed to settle for 3 hours prior to reactivation. Cells were fixed with 4% PFA for 15 minutes and permeabilised with PBS + 1% triton for 15 minutes. All further incubation steps occurred at 37 °C. Coverslips were blocked for 1 hour in PBS and 1% BSA, followed by 1 hour incubation in the appropriate primary antibody and 1 hour in Alexa-fluor conjugated secondary antibody (Invitrogen, 1/500) (70). Coverslips were mounted onto slides using Vectashield Mounting Medium with DAPI (Vector laboratories). Images were obtained using Zeiss LSM880 Inverted Confocal Microscope and processed using Zen Blue Software(71). For FISH, cells were seeded out and visualised per IF protocols whilst FISH was performed using ViewRNA Cell Plus Assay (ThermoFisher).

### Fluorescence recovery after photobleaching (FRAP)

TREx-BCBL1-RTA cells overexpressing GFP-SFPQ were used in FRAP experiments, cells were seeded onto poly-L-lysine-treated glass bottom dishes. Bleaching occurred using 488 nm laser at 100% intensity. Fluorescent intensity was measured pre-bleach and every 5 seconds for 210 seconds. A non-bleached area was measured concurrently as a control.

### Stimulated emission depletion microscopy (STED)

TREx-BCBL1-RTA cells were seeded onto washed poly-lysine treated cover slips and reactivated 3 hours later. Cells were fixed, permeabilised and blocked as per IF. Primary antibodies were used at twice the IF concentration and incubated for an hour at 37 °C. Secondary antibodies were Abberior Star Red (Abberior) and Abberior STAR 580 (Sigma Aldrich) used at 1/100 at 37 °C. Coverslips were mounted in Prolong Gold (ThermoFisher) and visualised on an Axio Observer microscope (Zeiss) with STEDYCON module (Abberior). STED images were deconvolved using Huygens Software (Scientific Volume Imaging).

### Viral reinfection and viral load assays

TREx-BCBL1-RTA cells were induced for 72 hours, DNA was extracted from cells and viral genomes measured via qPCR whilst the supernatant was added in a 1:1 ratio with naïve HEK-293Ts cell in DMEM. 24 hours after supernatant was added, RNA was harvested for analysis via qPCR.

### RNA immunoprecipitations

SFPQ and FLAG RIPs were performed in TREx-BCBL-1 RTA cells using EZ-Magna RIP RNA binding Immunoprecipitation Kit (Merck Millipore) as per manufacturer’s instructions. RNA was extracted and purified using TRIzol LS (Invitrogen) as per manufacturer’s instructions before analysis via qPCR. Samples were analysed using fold enrichment over % inputs.

### Immunoprecipitations

SFPQ IPs were performed in TREx-BCBL-1 RTA cells. SFPQ antibody was bound to either Protein A or Protein G magnetic beads (ThermoFisher) dependent on antibody species. Cell lysates were incubated overnight at 4 °C with the beads followed by washes. IPs were analysed alongside inputs via western blotting.

FLAG-Trap immunoprecipitations were performed in TREx-BCBL-1 RTA FLAG-ORF11 O/E cells as per the manufacturer’s protocol (Chromotek). Briefly, pre-washed FLAG-TRAP agarose beads (25 µL per sample) were combined with cell lysates, followed by a 2 hour incubation at 4 °C with rotation. The beads were washed three times, and samples were analysed alongside inputs via western blotting.

### Neutral comet assay

Comet assays were performed as published (72) using standard TREx-BCBL1-RTA cells, SFPQ depleted cells and SFPQ OE cells. Cells were suspended in low-melt agarose and added to microscope slides. Slides were incubated overnight at 37 °C in lysis buffer followed by electrophoresis and staining in 2.5 µg/ml of propidium iodide before visualisation on Zeiss LSM880 Inverted Confocal Microscope. Comet assays were analysed using CometScore2.0.

### Tandem-mass tagging coupled to liquid chromatography mass spectrometry (TMT LC-MS/MS)

Immunoprecipitation samples for quantitative MS analysis were sent to Bristol proteomics for TMT-LC/MS. Raw data files were obtained from Dr Kate Heesom (University of Bristol proteomics) and two metrics were derived: abundance change and abundance ratio. The former resulted from control values being subtracted from the sample values, whilst the latter resulted from sample values being divided by control values. To find true interactions percentage cut-offs relative to the bait protein were used: ≥5% abundance change, ≥10 abundance ratio. The proteins that met these criteria in both repeats were plotted on STRING to generate interaction networks for each pulldown. STRING website: https://string-db.org/cgi/input?sessionId=bPbwOPnnK8LL&input_page_show_search=on

### Statistical analysis

Except otherwise stated, graphical data shown represent mean ± standard deviation of mean (SD) using three or more biologically independent experiments. Differences between means were analysed by unpaired Student’s t-test. One-way Anova test was used for multiple comparisons. Statistics was considered significant at *P* < 0.05, with \**P* < 0.05, \*\**P* < 0.01 and \*\*\**P* < 0.001.

## Data availability

Source data for TMT-quantitative mass spectrometry have been deposited to the ProteomeXchange Consortium via the PRIDE partner repository with the dataset identifier PXD048834.

## Acknowledgements

We thank Professor Jae Jung (UCLA) for the TREx BCBL1-Rta cell line, Dr Jeffery Vieira (University of Washington) for the HEK-293T-rKSHV.219, Professors John Sinclair and Stacey Efstathiou (University of Cambridge) for HCMV (Merlin) and HSV-1 virus (SC16) stocks and Dr Edwin Chen (University of Westminster) for the psPAX2 and pVSV.G plasmids. We would like to thank Dr. Kate Heesom (Proteomics Facility, University of Bristol, UK) for the proteomics technical service and bioinformatics support. This work was supported in parts by the MRC (MR/X000060/1), White Rose BBSRC Doctoral Training Partnership in Mechanistic Biology (95519935) and University of Leeds Mary & Alice Smith Endowed Research Scholarship.

## Author Contributions

Conceptualization (KLH, EH, AW); Data curation (KLH, EH, WW); Formal Analysis (KLH, EH, REW, AW); Funding acquisition (AW); Investigation (KLH, EH, WW); Writing–original draft (KLH, EH, AW); Writing–review & editing (All authors).

## References

1. Wang, B., Zhang, L., Dai, T., Qin, Z., Lu, H., Zhang, L. and Zhou, F. Liquid-liquid phase separation in human health and diseases. Signal Transduct Target Ther. 2021, 6(1), p.290.

2. Fox, A.H., Lam, Y.W., Leung, A.K., Lyon, C.E., Andersen, J., Mann, M. and Lamond, A.I. Paraspeckles: a novel nuclear domain. Curr Biol. 2002, 12(1), pp.13–25.

3. Takakuwa, H., Yamazaki, T., Souquere, S., Adachi, S., Yoshino, H., Fujiwara, N., Yamamoto, T., Natsume, T., Nakagawa, S., Pierron, G. and Hirose, T. Shell protein composition specified by the lncRNA NEAT1 domains dictates the formation of paraspeckles as distinct membraneless organelles. Nat Cell Biol. 2023, 25(11), pp.1664–1675.

4. Naganuma, T. and Hirose, T. Paraspeckle formation during the biogenesis of long non-coding RNAs. RNA Biol. 2013, 10(3), pp.456–461.

5. Fox, A.H., Nakagawa, S., Hirose, T. and Bond, C.S. Paraspeckles: Where Long Noncoding RNA Meets Phase Separation. Trends Biochem Sci. 2018, 43(2), pp.124–135.

6. West, J.A., Mito, M., Kurosaka, S., Takumi, T., Tanegashima, C., Chujo, T., Yanaka, K., Kingston, R.E., Hirose, T., Bond, C., Fox, A. and Nakagawa, S. Structural, super-resolution microscopy analysis of paraspeckle nuclear body organization. J Cell Biol. 2016, 214(7), pp.817–830.

7. Hirose, T., Virnicchi, G., Tanigawa, A., Naganuma, T., Li, R., Kimura, H., Yokoi, T., Nakagawa, S., Benard, M., Fox, A.H. and Pierron, G. NEAT1 long noncoding RNA regulates transcription via protein sequestration within subnuclear bodies. Mol Biol Cell. 2014, 25(1), pp.169–183.

8. Fox, A.H. and Lamond, A.I. Paraspeckles. Cold Spring Harb Perspect Biol. 2010, 2(7), p.a000687.

9. An, H., Tan, J.T. and Shelkovnikova, T.A. Stress granules regulate stress-induced paraspeckle assembly. J Cell Biol. 2019, 218(12), pp.4127–4140.

10. Prasanth, K.V., Prasanth, S.G., Xuan, Z., Hearn, S., Freier, S.M., Bennett, C.F., Zhang, M.Q. and Spector, D.L. Regulating gene expression through RNA nuclear retention. Cell. 2005, 123(2), pp.249–263.

11. Imamura, K., Imamachi, N., Akizuki, G., Kumakura, M., Kawaguchi, A., Nagata, K., Kato, A., Kawaguchi, Y., Sato, H., Yoneda, M., Kai, C., Yada, T., Suzuki, Y., Yamada, T., Ozawa, T., Kaneki, K., Inoue, T., Kobayashi, M., Kodama, T., Wada, Y., Sekimizu, K. and Akimitsu, N. Long noncoding RNA NEAT1-dependent SFPQ relocation from promoter region to paraspeckle mediates IL8 expression upon immune stimuli. Mol Cell. 2014, 53(3), pp.393–406.

12. Stagsted, L.V.W., O’Leary, E.T., Ebbesen, K.K. and Hansen, T.B. The RNA-binding protein SFPQ preserves long-intron splicing and regulates circRNA biogenesis in mammals. Elife. 2021, 10.

13. Dittmer, D.P. and Damania, B. Kaposi sarcoma-associated herpesvirus: immunobiology, oncogenesis, and therapy. J Clin Invest. 2016, 126(9), pp.3165–3175.

14. Cesarman, E., Damania, B., Krown, S.E., Martin, J., Bower, M. and Whitby, D. Kaposi sarcoma. Nat Rev Dis Primers. 2019, 5(1), p.9.

15. Wen, K.W. and Damania, B. Kaposi sarcoma-associated herpesvirus (KSHV): molecular biology and oncogenesis. Cancer Lett. 2010, 289(2), pp.140–150.

16. Ganem, D. KSHV and the pathogenesis of Kaposi sarcoma: listening to human biology and medicine. J Clin Invest. 2010, 120(4), pp.939–949.

17. He, M., Cheng, F., da Silva, S.R., Tan, B., Sorel, O., Gruffaz, M., Li, T. and Gao, S.J. Molecular Biology of KSHV in Relation to HIV/AIDS-Associated Oncogenesis. Cancer Treat Res. 2019, 177, pp.23–62.

18. Dittmer, D., Lagunoff, M., Renne, R., Staskus, K., Haase, A. and Ganem, D. A cluster of latently expressed genes in Kaposi’s sarcoma-associated herpesvirus. J Virol. 1998, 72(10), pp.8309–8315.

19. Staskus, K.A., Zhong, W., Gebhard, K., Herndier, B., Wang, H., Renne, R., Beneke, J., Pudney, J., Anderson, D.J., Ganem, D. and Haase, A.T. Kaposi’s sarcoma-associated herpesvirus gene expression in endothelial (spindle) tumor cells. J Virol. 1997, 71(1), pp.715–719.

20. McClure, L.V. and Sullivan, C.S. Kaposi’s sarcoma herpes virus taps into a host microRNA regulatory network. Cell Host Microbe. 2008, 3(1), pp.1–3.

21. Broussard, G. and Damania, B. Regulation of KSHV Latency and Lytic Reactivation. Viruses. 2020, 12(9).

22. Arias, C., Weisburd, B., Stern-Ginossar, N., Mercier, A., Madrid, A.S., Bellare, P., Holdorf, M., Weissman, J.S. and Ganem, D. KSHV 2.0: a comprehensive annotation of the Kaposi’s sarcoma-associated herpesvirus genome using next-generation sequencing reveals novel genomic and functional features. PLoS Pathog. 2014, 10(1), p.e1003847.

23. Nicholas, J. Human herpesvirus 8-encoded proteins with potential roles in virus-associated neoplasia. Front Biosci. 2007, 12, pp.265–281.

24. Jackson, B.R., Noerenberg, M. and Whitehouse, A. A Novel Mechanism Inducing Genome Instability in Kaposi’s Sarcoma-Associated Herpesvirus Infected Cells. PLoS Pathog. 2014, 10(5), p.e1004098.

25. Grundhoff, A. and Ganem, D. Inefficient establishment of KSHV latency suggests an additional role for continued lytic replication in Kaposi sarcoma pathogenesis. J Clin Invest. 2004, 113(1), pp.124–136.

26. Giffin, L., Yan, F., Ben Major, M. and Damania, B. Modulation of Kaposi’s sarcoma-associated herpesvirus interleukin-6 function by hypoxia-upregulated protein 1. J Virol. 2014, 88(16), pp.9429–9441.

27. Manners, O., Murphy, J.C., Coleman, A., Hughes, D.J. and Whitehouse, A. Contribution of the KSHV and EBV lytic cycles to tumourigenesis. Curr Opin Virol. 2018, 32, pp.60–70.

28. Hughes, D.J., Wood, J.J., Jackson, B.R., Baquero-Perez, B. and Whitehouse, A. NEDDylation is essential for Kaposi’s sarcoma-associated herpesvirus latency and lytic reactivation and represents a novel anti-KSHV target. PLoS Pathog. 2015, 11(3), p.e1004771.

29. Baquero-Perez, B. and Whitehouse, A. Hsp70 Isoforms Are Essential for the Formation of Kaposi’s Sarcoma-Associated Herpesvirus Replication and Transcription Compartments. PLoS Pathog. 2015, 11(11), p.e1005274.

30. Vallery, T.K., Withers, J.B., Andoh, J.A. and Steitz, J.A. Kaposi’s Sarcoma-Associated Herpesvirus mRNA Accumulation in Nuclear Foci Is Influenced by Viral DNA Replication and Viral Noncoding Polyadenylated Nuclear RNA. J Virol. 2018, 92(13).

31. McCluggage, F. and Fox, A.H. Paraspeckle nuclear condensates: Global sensors of cell stress? Bioessays. 2021, 43(5), p.e2000245.

32. Sharma, N.R., Majerciak, V., Kruhlak, M.J. and Zheng, Z.M. KSHV inhibits stress granule formation by viral ORF57 blocking PKR activation. PLoS Pathog. 2017, 13(10), p.e1006677.

33. Hirose, T. and Nakagawa, S. Paraspeckles: possible nuclear hubs by the RNA for the RNA. Biomol Concepts. 2012, 3(5), pp.415–428.

34. Chen, L.L. and Carmichael, G.G. Altered nuclear retention of mRNAs containing inverted repeats in human embryonic stem cells: functional role of a nuclear noncoding RNA. Mol Cell. 2009, 35(4), pp.467–478.

35. Lellahi, S.M., Rosenlund, I.A., Hedberg, A., Kiaer, L.T., Mikkola, I., Knutsen, E. and Perander, M. The long noncoding RNA NEAT1 and nuclear paraspeckles are up-regulated by the transcription factor HSF1 in the heat shock response. J Biol Chem. 2018, 293(49), pp.18965–18976.

36. Knott, G.J., Bond, C.S. and Fox, A.H. The DBHS proteins SFPQ, NONO and PSPC1: a multipurpose molecular scaffold. Nucleic Acids Res. 2016, 44(9), pp.3989–4004.

37. Laurenzi, T., Palazzolo, L., Taiana, E., Saporiti, S., Ben Mariem, O., Guerrini, U., Neri, A. and Eberini, I. Molecular Modelling of NONO and SFPQ Dimerization Process and RNA Recognition Mechanism. Int J Mol Sci. 2022, 23(14).

38. Geiger, F., Acker, J., Papa, G., Wang, X., Arter, W.E., Saar, K.L., Erkamp, N.A., Qi, R., Bravo, J.P., Strauss, S., Krainer, G., Burrone, O.R., Jungmann, R., Knowles, T.P., Engelke, H. and Borodavka, Liquid-liquid phase separation underpins the formation of replication factories in rotaviruses. EMBO J. 2021, 40(21), p.e107711.

39. Murphy, J.C., Harrington, E.M., Schumann, S., Vasconcelos, E.J.R., Mottram, T.J., Harper, K.L., Aspden, J.L. and Whitehouse, A. Kaposi’s sarcoma-associated herpesvirus induces specialised ribosomes to efficiently translate viral lytic mRNAs. Nat Commun. 2023, 14(1), p.300.

40. Nakagawa, S., Yamazaki, T. and Hirose, T. Molecular dissection of nuclear paraspeckles: towards understanding the emerging world of the RNP milieu. Open Biol. 2018, 8(10).

41. Tang, S. and Zheng, Z.M. Kaposi’s sarcoma-associated herpesvirus K8 exon 3 contains three 5’-splice sites and harbors a K8.1 transcription start site. J Biol Chem. 2002, 277(17), pp.14547–14556.

42. Yamanegi, K., Tang, S. and Zheng, Z.M. Kaposi’s sarcoma-associated herpesvirus K8beta is derived from a spliced intermediate of K8 pre-mRNA and antagonizes K8alpha (K-bZIP) to induce p21 and p53 and blocks K8alpha-CDK2 interaction. J Virol. 2005, 79(22), pp.14207–14221.

43. Majerciak, V., Lu, M., Li, X. and Zheng, Z.M. Attenuation of the suppressive activity of cellular splicing factor SRSF3 by Kaposi sarcoma-associated herpesvirus ORF57 protein is required for RNA splicing. RNA. 2014, 20(11), pp.1747–1758.

44. Tagawa, T., Gao, S., Koparde, V.N., Gonzalez, M., Spouge, J.L., Serquina, A.P., Lurain, K., Ramaswami, R., Uldrick, T.S., Yarchoan, R. and Ziegelbauer, J.M. Discovery of Kaposi’s sarcoma herpesvirus-encoded circular RNAs and a human antiviral circular RNA. Proc Natl Acad Sci U S A. 2018.

45. Harper, K.L., Mottram, T.J., Anene, C.A., Foster, B., Patterson, M.R., McDonnell, E., Macdonald, A., Westhead, D. and Whitehouse, A. Dysregulation of the miR-30c/DLL4 axis by circHIPK3 is essential for KSHV lytic replication. EMBO Rep. 2022, 23(5), p.e54117.

46. Abere, B., Li, J., Zhou, H., Toptan, T., Moore, P.S. and Chang, Y. Kaposi’s Sarcoma-Associated Herpesvirus-Encoded circRNAs Are Expressed in Infected Tumor Tissues and Are Incorporated into Virions. mBio. 2020, 11(1).

47. Tagawa, T., Koparde, V.N. and Ziegelbauer, J.M. Identifying and characterizing virus-encoded circular RNAs. Methods. 2021.

48. Wongwiwat, W., Fournier, B., Bassano, I., Bayoumy, A., Elgueta Karstegl, C., Styles, C., Bridges, R., Lenoir, C., BoutBoul, D., Moshous, D., Neven, B., Kanda, T., Morgan, R.G., White, R.E., Latour, S. and Farrell, P.J. Epstein-Barr Virus Genome Deletions in Epstein-Barr Virus-Positive T/NK Cell Lymphoproliferative Diseases. J Virol. 2022, 96(12), p.e0039422.

49. Neuhierl, B., Feederle, R., Hammerschmidt, W. and Delecluse, H.J. Glycoprotein gp110 of Epstein-Barr virus determines viral tropism and efficiency of infection. Proc Natl Acad Sci U S A. 2002, 99(23), pp.15036–15041.

50. Cavallin, L.E., Goldschmidt-Clermont, P. and Mesri, E.A. Molecular and cellular mechanisms of KSHV oncogenesis of Kaposi’s sarcoma associated with HIV/AIDS. PLoS Pathog. 2014, 10(7), p.e1004154.

51. Rosemarie, Q. and Sugden, B. Epstein-Barr Virus: How Its Lytic Phase Contributes to Oncogenesis. Microorganisms. 2020, 8(11).

52. Bladen, C.L., Udayakumar, D., Takeda, Y. and Dynan, W.S. Identification of the polypyrimidine tract binding protein-associated splicing factor.p54(nrb) complex as a candidate DNA double-strand break rejoining factor. J Biol Chem. 2005, 280(7), pp.5205–5210.

53. Udayakumar, D. and Dynan, W.S. Characterization of DNA binding and pairing activities associated with the native SFPQ.NONO DNA repair protein complex. Biochem Biophys Res Commun. 2015, 463(4), pp.473–478.

54. Jaafar, L., Li, Z., Li, S. and Dynan, W.S. SFPQ*NONO and XLF function separately and together to promote DNA double-strand break repair via canonical nonhomologous end joining. Nucleic Acids Res. 2017, 45(4), pp.1848–1859.

55. Boija, A., Klein, I.A. and Young, R.A. Biomolecular Condensates and Cancer. Cancer Cell. 2021, 39(2), pp.174–192.

56. Gordon, P.M., Hamid, F., Makeyev, E.V. and Houart, C. A conserved role for the ALS-linked splicing factor SFPQ in repression of pathogenic cryptic last exons. Nat Commun. 2021, 12(1), p.1918.

57. Vendruscolo, M. and Fuxreiter, M. Protein condensation diseases: therapeutic opportunities. Nat Commun. 2022, 13(1), p.5550.

58. Nam, J. and Gwon, Y. Neuronal biomolecular condensates and their implications in neurodegenerative diseases. Front Aging Neurosci. 2023, 15, p.1145420.

59. Brownsword, M.J. and Locker, N. A little less aggregation a little more replication: Viral manipulation of stress granules. Wiley Interdiscip Rev RNA. 2023, 14(1), p.e1741.

60. Naganuma, T., Nakagawa, S., Tanigawa, A., Sasaki, Y.F., Goshima, N. and Hirose, T. Alternative 3’-end processing of long noncoding RNA initiates construction of nuclear paraspeckles. EMBO J. 2012, 31(20), pp.4020–4034.

61. Hennig, S., Kong, G., Mannen, T., Sadowska, A., Kobelke, S., Blythe, A., Knott, G.J., Iyer, K.S., Ho, D., Newcombe, E.A., Hosoki, K., Goshima, N., Kawaguchi, T., Hatters, D., Trinkle-Mulcahy, L., Hirose, T., Bond, C.S. and Fox, A.H. Prion-like domains in RNA binding proteins are essential for building subnuclear paraspeckles. J Cell Biol. 2015, 210(4), pp.529–539.

62. Dattilo, D., Di Timoteo, G., Setti, A., Giuliani, A., Peruzzi, G., Beltran Nebot, M., Centron-Broco, A., Mariani, D., Mozzetta, C. and Bozzoni, I. The m(6)A reader YTHDC1 and the RNA helicase DDX5 control the production of rhabdomyosarcoma-enriched circRNAs. Nat Commun. 2023, 14(1), p.1898.

63. Li, Y., Chen, B., Zhao, J., Li, Q., Chen, S., Guo, T., Li, Y., Lai, H., Chen, Z., Meng, Z., Guo, W., He, X. and Huang, S. HNRNPL Circularizes ARHGAP35 to Produce an Oncogenic Protein. Adv Sci (Weinh). 2021, 8(13), p.2001701.

64. Takeuchi, A., Iida, K., Tsubota, T., Hosokawa, M., Denawa, M., Brown, J.B., Ninomiya, K., Ito, M., Kimura, H., Abe, T., Kiyonari, H., Ohno, K. and Hagiwara, M. Loss of Sfpq Causes Long-Gene Transcriptopathy in the Brain. Cell Rep. 2018, 23(5), pp.1326–1341.

65. Roundtree, I.A., Luo, G.Z., Zhang, Z., Wang, X., Zhou, T., Cui, Y., Sha, J., Huang, X., Guerrero, L., Xie, P., He, E., Shen, B. and He, C. YTHDC1 mediates nuclear export of N(6)-methyladenosine methylated mRNAs. Elife. 2017, 6.

66. Cheng, Y., Xie, W., Pickering, B.F., Chu, K.L., Savino, A.M., Yang, X., Luo, H., Nguyen, D.T., Mo, S., Barin, E., Velleca, A., Rohwetter, T.M., Patel, D.J., Jaffrey, S.R. and Kharas, M.G. N(6)-Methyladenosine on mRNA facilitates a phase-separated nuclear body that suppresses myeloid leukemic differentiation. Cancer Cell. 2021, 39(7), pp.958–972 e958.

67. Ye, F., Chen, E.R. and Nilsen, T.W. Kaposi’s Sarcoma-Associated Herpesvirus Utilizes and Manipulates RNA N(6)-Adenosine Methylation To Promote Lytic Replication. J Virol. 2017, 91(16).

68. Hollingworth, R., Skalka, G.L., Stewart, G.S., Hislop, A.D., Blackbourn, D.J. and Grand, R.J. Activation of DNA Damage Response Pathways during Lytic Replication of KSHV. Viruses. 2015, 7(6), pp.2908–2927.

69. Baquero-Perez, B., Antanaviciute, A., Yonchev, I.D., Carr, I.M., Wilson, S.A. and Whitehouse, A. The Tudor SND1 protein is an m(6)A RNA reader essential for replication of Kaposi’s sarcoma-associated herpesvirus. Elife. 2019, 8.

70. Goodwin, D.J., Hall, K.T., Giles, M.S., Calderwood, M.A., Markham, A.F. and Whitehouse, A. The carboxy terminus of the herpesvirus saimiri ORF 57 gene contains domains that are required for transactivation and transrepression. Journal of General Virology. 2000, 81(9), pp.2253–2658.

71. Manners, O., Baquero-Perez, B., Mottram, T.J., Yonchev, I.D., Trevelyan, C.J., Harper, K.L., Menezes, S., Patterson, M.R., Macdonald, A., Wilson, S.A., Aspden, J.L. and Whitehouse, A. m(6)A Regulates the Stability of Cellular Transcripts Required for Efficient KSHV Lytic Replication. Viruses. 2023, 15(6).

72. Olive, P.L. and Banath, J.P. The comet assay: a method to measure DNA damage in individual cells. Nat Protoc. 2006, 1(1), pp.23–29.

