## Supplementary figures for "Virus-induced paraspeckle-like condensates are essential hubs for gene expression and their formation drives genomic instability"

Supplementary Figures 1-6

Supplementary Table 1

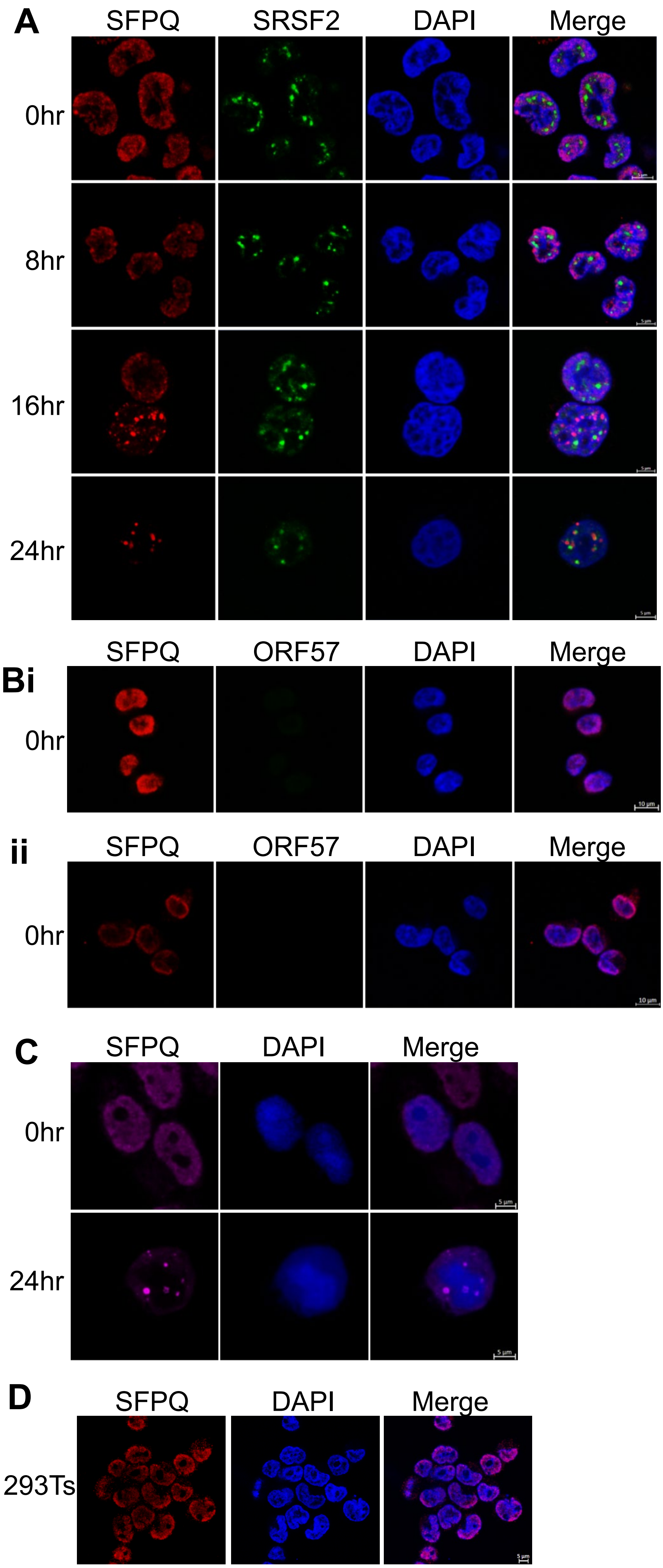


**Supplementary Figure 1:**

**(A).** IF of TREx-BCBL1-Rta cells at 0, 8, 16 and 24 hours staining for SFPQ and SRSF2. (**B**)**.** IF of TREx-BCBL1-Rta cells during latency, with a 24 hour pre-treatment serum starvation (i) or TREx-BCBL1-RTA cells exposed to osmotic stress (ii). Stained for SFPQ and ORF57. **(C).** IF of HEK-293T-rKSHV.219 cells at 0 and 24 hours post-reactivation with antibodies against SFPQ (red). **(D).** IF of HEK-293T cells with antibodies against SFPQ (red). Scale bars are 5 µm in A-D and 10 µm in B.

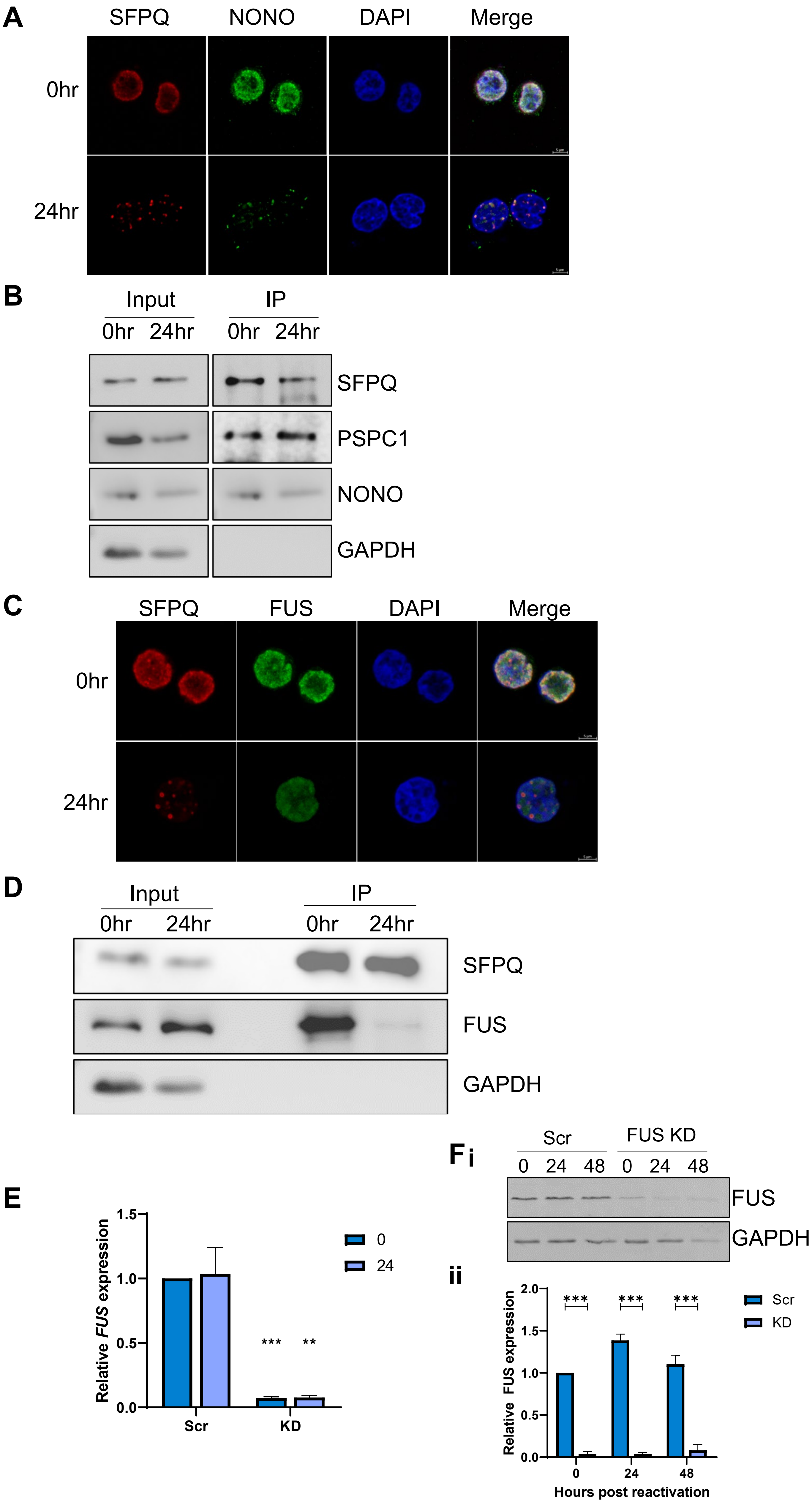
A
SFPQ
NONO
DAPI
Merge
0hr
24hr
B
Input
IP
0hr
24hr
0hr
24hr
SFPQ
PSPC1
NONO
GAPDH
C
SFPQ
FUS
DAPI
Merge
0hr
24hr
D
Input
IP
0hr
24hr
0hr
24hr
SFPQ
FUS
GAPDH
E
Relative FUS expression
1.5
1.0
0.5
0.0
0
24
***
**
Scr
KD
Fi
Scr
FUS KD
0
24
48
0
24
48
FUS
GAPDH
ii
Relative FUS expression
2.0
1.5
1.0
0.5
0.0
***
***
***
Scr
KD
0
24
48
Hours post reactivation

G

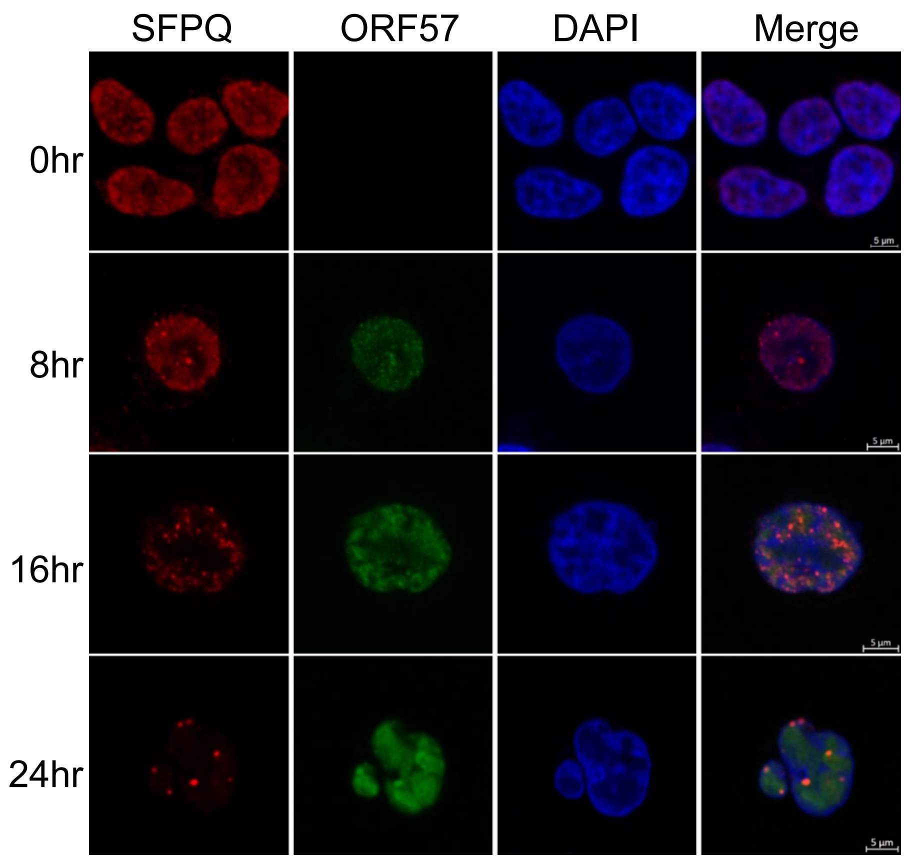


H

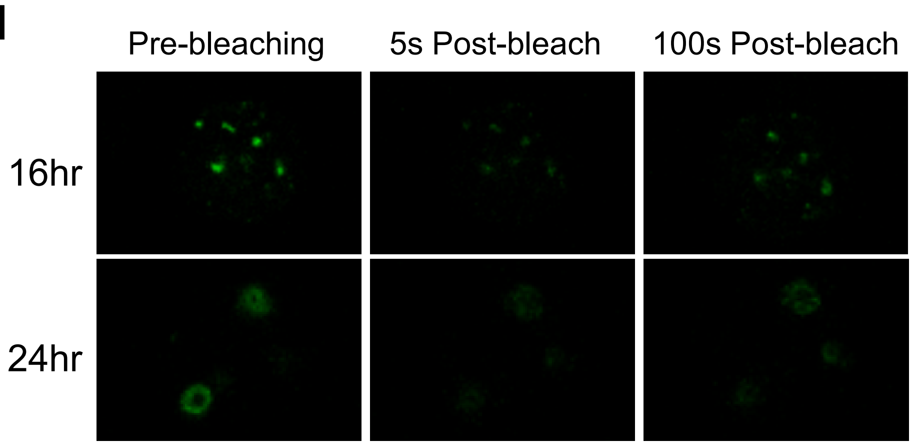


**Supplementary Figure 2:**

**(A)** IF of TREx at 0 and 24 hours with staining for SFPQ (red) and NONO (green). **(B)**. Co-IP of SFPQ probed with antibodies against SFPQ, PSPC1, NONO and GAPDH (n=3). **(C).** IF of TREx at 0 and 24 hours with staining for SFPQ (red) and FUS (green). **(D)**. Representative western blot of SFPQ co-IP in TREx at 0 and 24 hours with antibodies against SFPQ, FUS and GAPDH. n=3. **(E).** qPCR of *FUS* in scr and FUS KD TREx cells at 0 and 24 hours. GAPDH was used a housekeeper, n=3. **(F).** Representative western blot for levels of FUS in scr and FUS KD cells at 0, 24 and 48 hours post-lytic induction (i). GAPDH was used as a loading control, densitometry analysis for latent and 48 hours post-lytic induction was performed on n=3 (ii). **(G)**. IF of TREx cells at 0, 8, 16 and 24 hours with SFPQ (red) and ORF57 (green) staining. **(H).** FRAP images of GFP-SFPQ O/E TREx cells at 16 and 24 hours post-reactivation. Images were taken 5 seconds pre-bleaching, 5 seconds post-bleaching and 100 seconds post-bleaching. Scale bars are 5 µm. All repeats are biological. In E-F data are presented as mean ± SD. Unpaired student T-test. *P < 0.05, **P < 0.01 and ***P < 0.001.

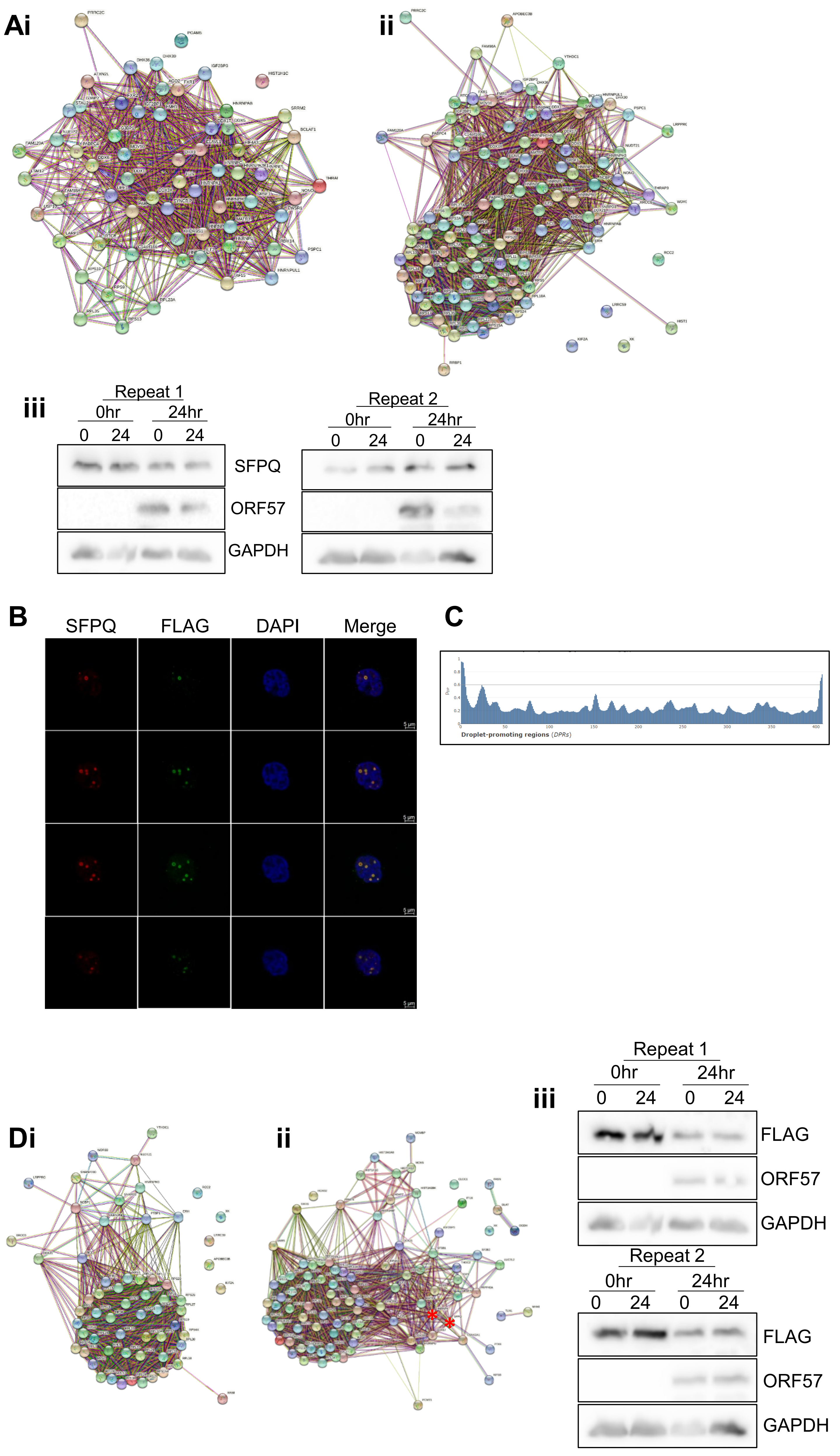
Ai
ii
iii
Repeat 1
0hr
24hr
0 24 0 24
SFPQ
ORF57
GAPDH
Repeat 2
0hr
24hr
0 24 0 24
B
SFPQ
FLAG
DAPI
Merge
5 µm
C
Droplet-promoting regions (DPRs)
Di
ii
iii
Repeat 1
0hr
24hr
0 24 0 24
FLAG
ORF57
GAPDH
Repeat 2
0hr
24hr
0 24 0 24
FLAG
ORF57
GAPDH

**E**

| DEAD/DEAH box helicases | hnRNPs | Other RNA processing factors |
|---|---|---|
| DDX1 | hnRNP A1 | ADAR |
| DDX17 | hnRNP A2B1 | APOBEC3B |
| DDX21 | hnRNP AB | ELAV1 |
| DDX3X | hnRNP C | ERH |
| DDX5 | hnRNP D | FAM98A |
| DHX30 | hnRNP DL | FBL |
| DHX36 | hnRNP H1 | FMR1 |
| DHX9 | hnRNP H3 | IGF2BP3 |
| | hnRNP K | LUC7L2 |
| | hnRNP L | MOV10 |
| | hnRNP M | NCL |
| | hnRNP Q | PABPC1 |
| | hnRNP R | PABPC4 |
| | hnRNP U | PRPF40A |
| | hnRNP UL1 | PTBP1 |
| | | RARS1 |
| | | RTCB |
| | | RTRAF |
| | | SF3BP |
| | | SNRNP200 |
| | | SNRPA |
| | | SNRPD3 |
| | | SRSF1 |
| | | UPF1 |
| | | VARS1 |
| | | WDR33 |
| | | YTHDC1 |

**F**

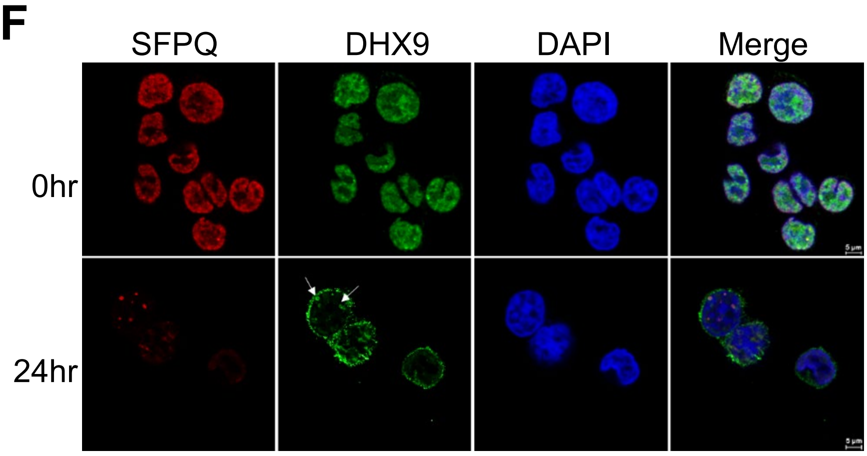


**G**

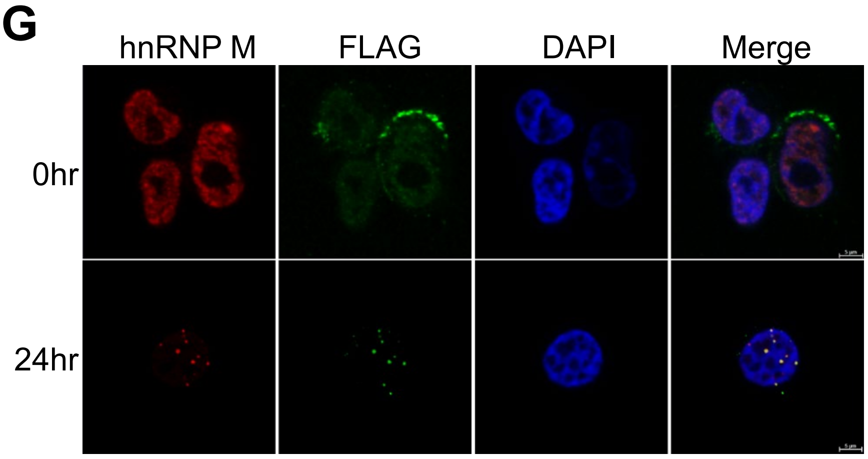

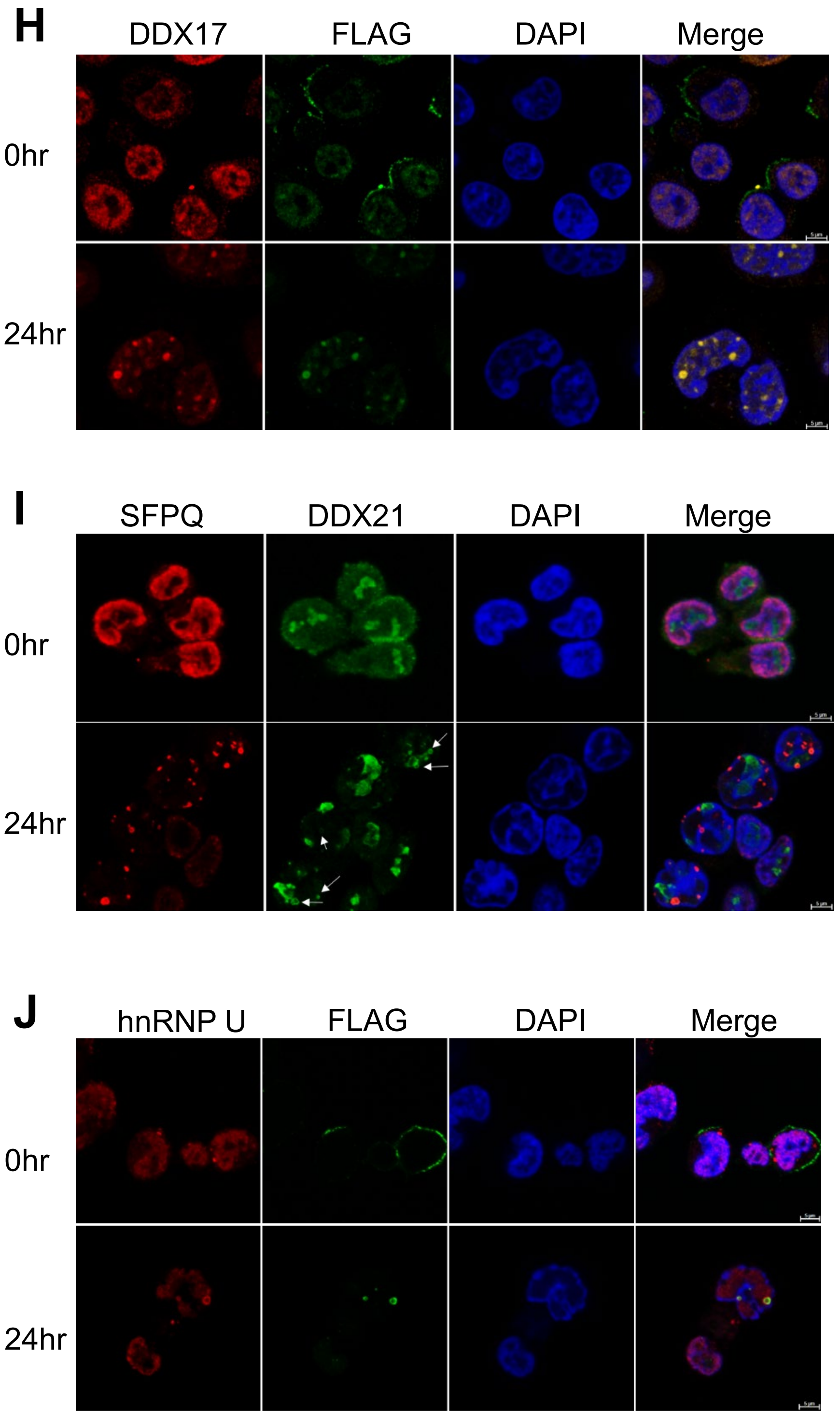


**Supplementary Fig 3:**

**(A).** String analysis of TMT LC-MS/MS of SFPQ Co-IPs in TREx-BCBL1-RTA cells at 0 (i) and 24 hours (ii). (n=2). Inputs were analysed via western blot to confirm presence of SFPQ and lytic replication (iii). **(B).** IF analysis taken through different z-planes of a cell stained for SFPQ (red) and FLAG (green) at 24 hours in TREx FLAG-ORF11 OE cells. **(C).** Protein disorder prediction for ORF11 using FuzDrop software. **(D).** String analysis of TMT LC-MS/MS of FLAG (ORF11) Co-IPs in TREx FLAG-ORF11 O/E cells. (n=2) at 0 (i) and 24 hours (ii). Inputs were analysed via western blot to confirm presence of FLAG-ORF11 and lytic replication (iii). * denotes proteins of interest **(E).** Table highlighting proteins of interest within the SFPQ or FLAG-ORF11 TMT-MS at 24 hours. Proteins are divided into DEAD/DEAH box helicases, hnRNPs or other known RNA processing factors. **(F-I)** IF analysis of some of the TMT-MS enriched targets at 0 and 24 hours in TREx or TREx FLAG-ORF11 O/E cells with staining for **(F)** SFPQ (red) and DHX9 (green), **(G)** hnRNP M (red) and FLAG (green), **(H)**, DDX17 (red) and FLAG (green), **(I)** SFPQ (red) and DDX21 (green) **(J)** hnRNP U (red) and FLAG (green). White arrows are used to highlight condensates. Scale bars are 5 µm in length.

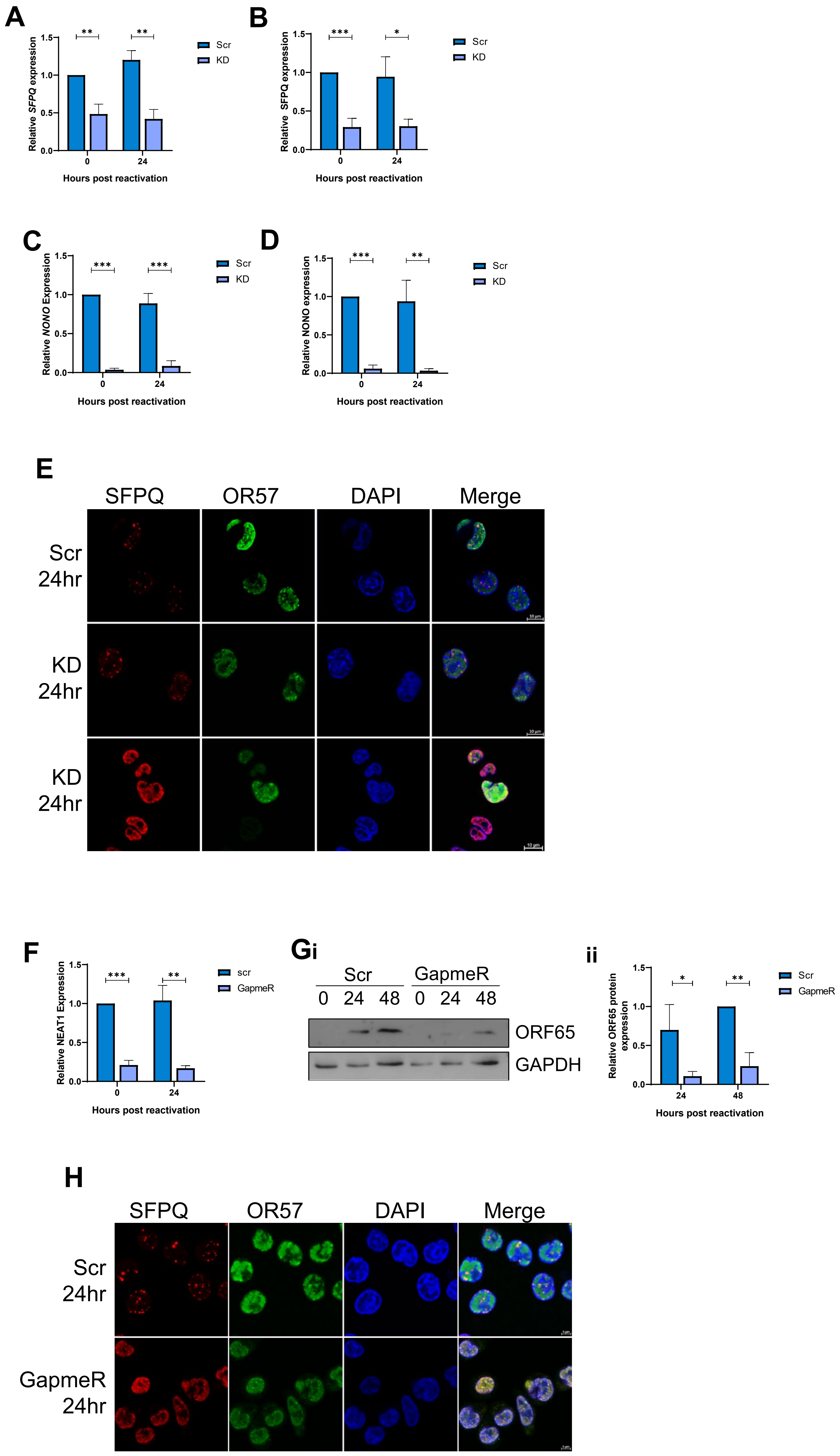
A
Relative SFPQ expression
1.5
1.0
0.5
0.0
**
0
24
Hours post reactivation
Scr
KD
B
Relative SFPQ expression
***
*
Hours post reactivation
Scr
KD
C
Relative NONO Expression
***
Hours post reactivation
Scr
KD
D
Relative NONO expression
***
**
Hours post reactivation
Scr
KD
E
SFPQ
OR57
DAPI
Merge
Scr 24hr
KD 24hr
KD 24hr
F
Relative NEAT1 Expression
***
**
Hours post reactivation
scr
GapmeR
Gi
Scr
GapmeR
0 24 48 0 24 48
ORF65
GAPDH
ii
Relative ORF65 protein expression
*
**
24
48
Hours post reactivation
Scr
GapmeR
H
SFPQ
OR57
DAPI
Merge
Scr 24hr
GapmeR 24hr

**Supplementary Figure 4.**

**(A).** qPCR analysis of *SFPQ* levels at 0 and 24 hours in TREx scr and SFPQ KD cells. GAPDH was used as a housekeeper (n=3). **(B).** Densitometry analysis of SFPQ protein levels in 0 and 24 hours in scr and SFPQ KD TREx cells (n=3). **(C).** qPCR analysis of *NONO* levels in scr and NONO KD TREx cells at 0 and 24 hours. GAPDH was used as a housekeeper. (n=3). **(D).** Densitometry analysis of NONO protein levels in 0 and 24 hours in scr and NONO KD TREx cells (n=3). **(E).** IF in scr or NONO KD TREx cells with staining against SFPQ (red) and ORF57 (green). **(F).** qPCR analysis of *NEAT1* levels in scr or GapmeR treated TREx cells at 0 or 24 hours. GAPDH was used as a housekeeper, n=3. **(G).** Representative western blot of ORF65 in scrambled and *NEAT1* GapmeR treated TREx at 0, 24 hours and 48 hours (i). GAPDH was used a loading control. Densitometry analysis on n=3 for ORF65 (ii). **(H).** IF analysis of scrambled and *NEAT1* GapmeR treated TREx at 24 hours with staining for SFPQ (red) and ORF57 (green). Scale bars are 10 µm in E and 5 µm in H. All repeats are biological. In A-D, F-G data are presented as mean ± SD. $^*P < 0.05$, $^{**}P < 0.01$ and $^{***}P < 0.001$ (unpaired Student's t-test).

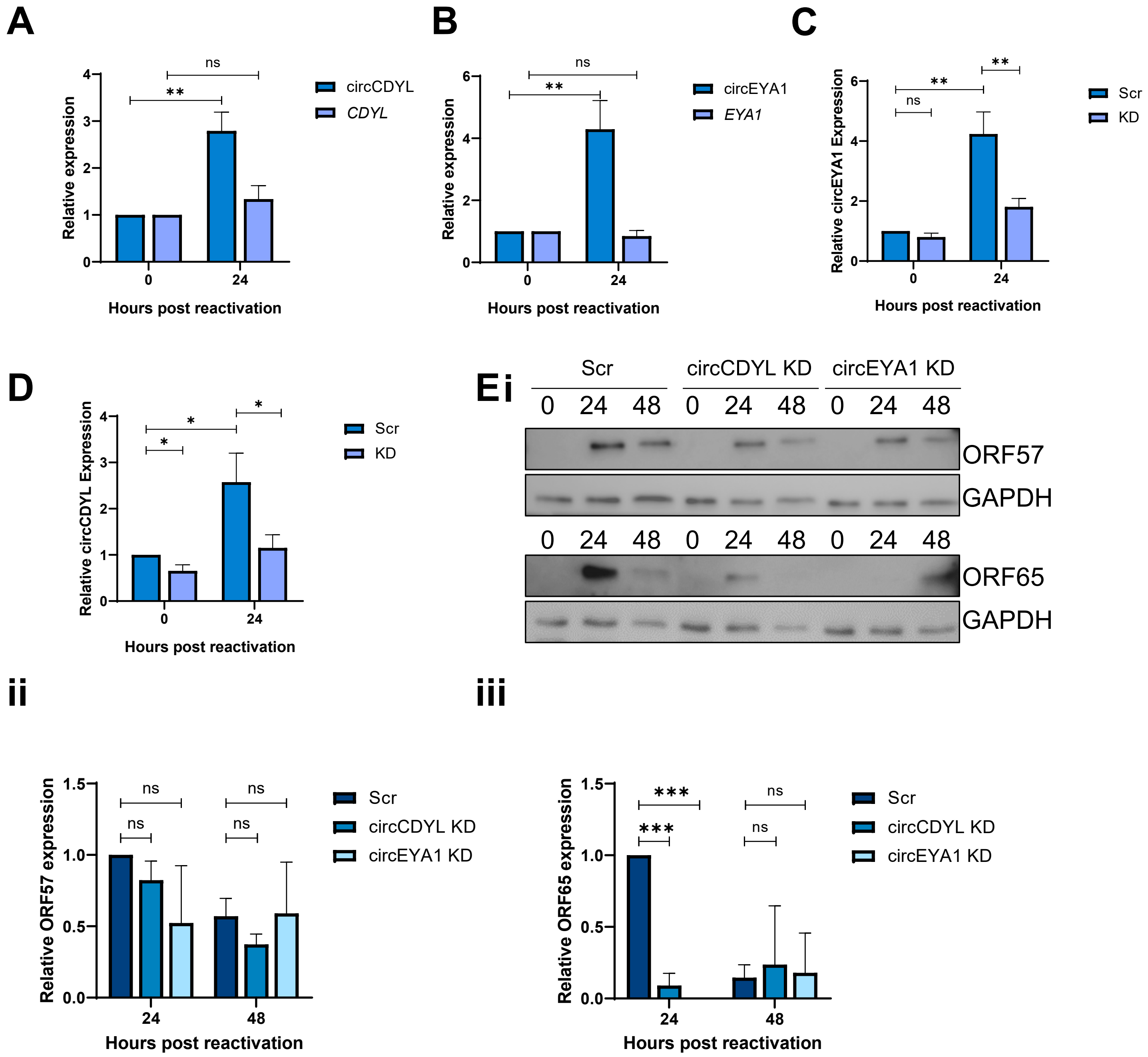


**Supplementary Figure 5.**

**(A).** qPCR of circCDYL and *CDYL* levels at 0 and 24 hours in TREx, GAPDH was used as a housekeeper, n=3. **(B).** qPCR of circEYA1 and *EYA1* levels at 0 and 24 hours in TREx cells, GAPDH was used as a housekeeper, n=3. **(C)** qPCR analysis of circEYA1 levels in scramble and circEYA1 KD TREx cells at 0 and 24 hours. GAPDH was used as a housekeeper, n=3. **(D)** qPCR analysis of circCDYL levels in scramble and circCDYL KD TREx cells at 0 and 24 hours. GAPDH was used as a housekeeper, n=3. **(E).** Representative western blot of ORF57 and ORF65 levels in scrambled, circCDYL KD and circEYA1 KD TREx cells at 0, 24 and 48 hours (i). GAPDH was used as a loading control. Densitometry analysis was performed on n=3 for ORF57 (ii) and ORF65 (iii). All repeats are biological. In A-E data are presented as mean ± SD. $^{*}P < 0.05$, $^{**}P < 0.01$ and $^{***}P < 0.001$. Unpaired Student's t-test was used for A-D and a one-way ANOVA was performed for E.

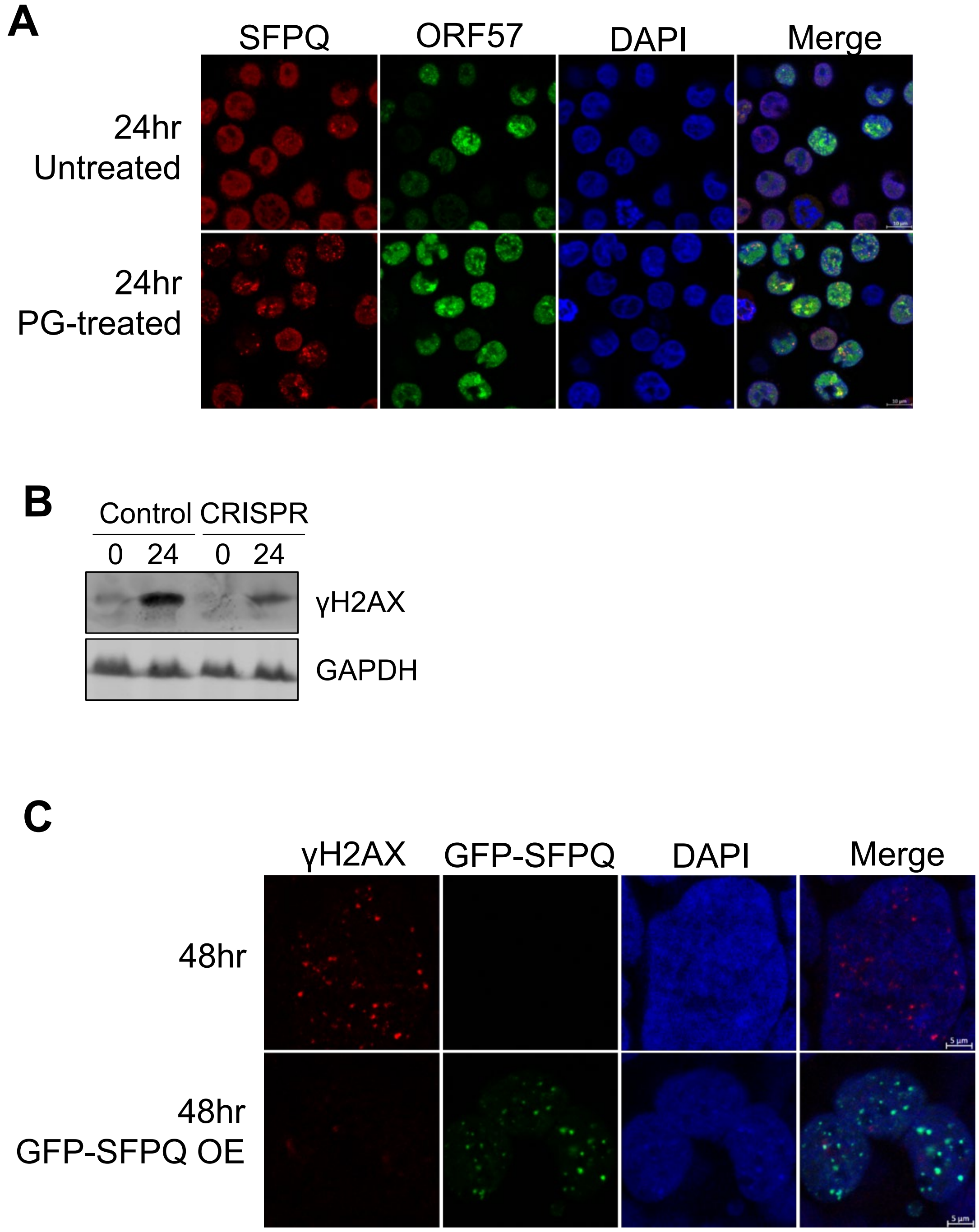


**Supplementary Figure 6.**

**(A).** IF of untreated or 8 hour PG treated TREx cells. Cells were reactivated for 24 hours, with PG added at 16 hours post-lytic induction. Antibodies against SFPQ (red) and ORF57 (green) were used. **(B)**. Representative western blot of γH2AX at 0 and 24 hours in control and ORF11 CRISPR cells. GAPDH was used as a loading control, n=3. **(C).** IF of rJJ-L3-#1 at 48 hours either positive or negative for GFP-SFPQ OE. Cells were stained for γH2AX.Scale bars are 10 μm in A and 5 μm in C. All repeats are biological.

**Supplementary Table 1: Primer sequences**

| Primer | Forward | Reverse |
|---|---|---|
| GAPDH | TGTCAGTGGTGGACCTGA | GTGGTCCTTGAGGGCAATG |
| ORF57 | GCCATAATCAAGCGTACTGG | GCAGAGAAATATTGCGGTGT |
| K8 | AGGACCACACATTTCGCAAC | ACCCCTTGTCAGTTCTTC |
| PAN | ATAGGCGACAAAGTGAGGTGGCAT | TAACATTGAAAGAGCGCTCCCAGC |
| ORF59 | GGTCCGGATATGCTCCTAGTT | CAGCATGCTCACGAGGAATA |
| ORF4 V1 | CGATTTGTGCACGGAAGA | GGAGTGTTGGTTCTCGC |
| ORF4 V2 | CGTTTGCATCCAACACCCAAT | TGTTGGTTCTCGCGGTCTC |
| ORF4 V3 | TCCTGCCAACATCCGAAGG | GAGGGAGTGTTGGTTCTCGC |
| ORF65 | AAGGTGAGAGACCCCGTGAT | TCCAGGGTATTCATGCGAGC |
| Ea-D | CTAGCCGTCCTGTCCAAGTGC | AGCCAAACGCTCCTTGCCCA |
| SFPQ | ACAGGGAAAGGCATTGTTGA | TCATCTAGTTGTTCAAGTGGTTCC |
| NONO | TGATGAAGAGGGACTTCCAGA | AGCGCATGGCATATTCATACT |
| NEAT1 | AGTACCCTGAGAGCCAGTATTGGT | GGCAGCTGAGTCAATCTCCTTT |
| circPAN | ACCAGACGGCAAGGTTTTA | TCGTTAGTCAACCTAGCAAAACA |
| circvIRF4 | CTCCGTGTGGATACCAGTGA | TGGTCCCACGCAACAGTCT |
| vIRF4 | CCCAACAGGCCAGCTACATAA | CTTCGTGGAACTCTGAGACGC |
| EYA1 | GAGCTGATGGCTCCGAGTTT | GCTATGCGGGCTGGTTAGAT |
| circEYA1 | TTGCTTACTGGGTCCTACGC | TACTGCTCCCAATTGCTGAA |
| CDYL | CGAGGAGCTGTACGAGGTTG | ACGAGTGCCTAAGGAGAGGT |
| circCDYL | ACCCACTAGTGCCTCAGGTG | CTCGCTGTCATGCCTTTCC |
| circSPECC1 | GAGAGCTGCGAAGTTCAAGA | GCCTGTCCGTTTAGTTGTTGT |
